# Robust W1282X-CFTR rescue by a small molecule GSPT1 degrader

**DOI:** 10.1101/2021.09.28.462149

**Authors:** Rhianna E. Lee, Catherine A. Lewis, Lihua He, Samuel C. Gallant, Teresa M. Mascenik, Hong Dang, Lisa C. Morton, John T. Minges, Jonathan W. Theile, Neil A. Castle, Michael R. Knowles, Adam J. Kimple, Scott H. Randell

**Author notes:** First author (Rhianna E. Lee). Contributed equally to this work (Catherine A. Lewis, Lihua He). Corresponding author, Scott Randell, 1117 Marsico Hall, Campus Box 7248, Chapel Hill, NC 27599-7248, /.

## Abstract

With the approval of elexacaftor/tezacaftor/ivacaftor (trade name Trikafta), the vast majority of people with cystic fibrosis (CF) are eligible for CF transmembrane conductance regulator (CFTR) modulator therapy. Remaining individuals have premature termination codons or rare *CFTR* variants with limited treatment options. Although clinical modulator response can be reliably predicted using primary airway epithelial cells, primary cells carrying rare *CFTR* variants are scarce. To overcome this obstacle, these cells can be expanded by overexpression of mouse *Bmi-1* and human *TERT* (hTERT). We therefore used this approach to develop two non-CF and three CF (F508del/F508del, F508del/S492F, W1282X/W1282X) nasal cell lines and two W1282X/W1282X bronchial cell lines. Bmi-1/hTERT cell lines recapitulated primary cell morphology and ion transport function. The F508del/F508del and F508del/S492F cell lines robustly responded to Trikafta, which was mirrored in the parent primary cells and the cell donors’ clinical response. CC-90009, a novel cereblon E3 ligase modulator targeting the GSPT1 protein, rescued ~20% of wildtype CFTR function in our panel of W1282X/W1282X cell lines and primary cells. Intriguingly, CC-90009 also diminished epithelial sodium channel function. These studies demonstrate that Bmi-1/hTERT cell lines faithfully mirror primary cell responses to CFTR modulators and illustrate novel therapeutic approaches for the W1282X CFTR variant.

## Introduction

Cystic fibrosis (CF) is a life-limiting genetic disease affecting approximately 70,000 people worldwide (1). Severe pathology develops in the lungs, where absent or dysfunctional CF transmembrane conductance regulator (CFTR) protein leads to the accumulation of thick airway mucus, impaired mucus transport, chronic infection and inflammation, and, eventually, bronchiectasis (2). Historically, treatments have been limited to symptom management. However, the 2012 US Food and Drug Administration (FDA) approval of the first small molecule CFTR modulator ivacaftor ushered in a new era of CF precision medicine (3). In contrast to previous treatment approaches, CFTR modulators treat the underlying cause of disease by directly acting on the CFTR protein to correct folding, trafficking, function, or stability. With additional CFTR modulator approvals (4–6) culminating in the 2019 approval of a triple combination therapy, elexacaftor/tezacaftor/ivacaftor (trade name Trikafta) (7), as many as 90% of people with CF are now eligible for an FDA-approved modulator therapy.

However, developing therapies for the remaining individuals has proven challenging. This is in part because this group harbors a wide range of rare *CFTR* variants. Indeed, more than 1,200 *CFTR* variants are carried by five or fewer people worldwide (8; CFTR2.org). For these individuals, a well-powered clinical trial is not possible. Thus, to extend life-changing treatment to all people with CF, the approach for evaluating candidate therapies must evolve.

The FDA set the precedent for such a change in 2017 when they expanded the use of ivacaftor to patient populations harboring one of twenty-three relatively rare *CFTR* variants (https://www.accessdata.fda.gov/scripts/cder/daf/index.cfm?event=overview.process&ApplNo=203188). Although drug label expansions are common, this instance was particularly groundbreaking because the FDA based their decision purely on *in vitro* data rather than a clinical trial (9). This new paradigm relies on the fidelity of *in vitro* systems to accurately predict a clinical response.

Primary CF human bronchial epithelial cells (HBECs) obtained at the time of lung transplantation have served as the gold standard for assessing CFTR rescue *in vitro* (10–12). However, the supply of CF explant lungs is limited, particularly for rare *CFTR* variants. HBECs can also be obtained by bronchial brushing, but the procedure is invasive and yields low cell numbers (13). A readily-available alternative to HBECs is primary human nasal epithelial cells (HNECs), which are increasingly being used as a model for the lower airways. HNECs can be obtained via nasal curettage, a non-surgical and well-tolerated method (14). A direct comparison of paired HBEC and HNEC samples demonstrated that mature airway cell markers and CFTR activity with and without modulator treatment are preserved (13). However, like bronchial brushing, nasal curettage yields a limited supply of cells.

Previous work by our lab and others has shown that expression of mouse B cell-specific Moloney murine leukemia virus integration site 1 (Bmi-1) and human telomerase reverse transcriptase (hTERT) enable robust expansion of bronchial epithelial cells that can be differentiated and assayed for CFTR function for up to 15 passages (15, 16). Other groups have applied this method to nasopharyngeal biopsies (17), but to our knowledge, this technique has not been used for nasal curettage samples. We hypothesized that Bmi-1 and hTERT expression would enhance HNEC growth properties while maintaining their ability to differentiate into a polarized pseudostratified epithelium. We propose that this method could be applied to rare genotype HNECs and HBECs as they become available to create the cellular resources required for personalized medicine and drug discovery.

Here, we created five nasal and two bronchial Bmi-1/hTERT cell lines that recapitulate primary cell morphology and ion transport function for at least 15 passages. By examining cell lines with F508del/F508del and F508del/S492F CFTR genotypes and comparing them to the parent primary cells, we assessed the fidelity of CFTR modulator responses and the percent of normal CFTR activity restored. We also compared the *in vitro* cell culture and *in vivo* cell donors’ clinical responses to Trikafta.

We then used this approach to evaluate a potential modulator therapy for patients homozygous for W1282X, a nonsense mutation that generates a premature termination codon (PTC) in the *CFTR* transcript, predisposing *CFTR* mRNA to nonsense-mediated decay (NMD) and absent or truncated protein. Without a targetable protein, modulator therapies are ineffective, leaving these patients without treatment options. Recent studies illustrate that the cereblon E3 ubiquitin ligase modulator, CC-90009, promotes PTC readthrough of nonsense mutations associated with inherited diseases (18). Using one nasal and two bronchial Bmi-1/hTERT W1282X/W128X CFTR cell lines, we tested CC-90009 effects, again showing fidelity between cell lines and primary cells and importantly both predicted and unanticipated electrophysiological responses that would be beneficial in CF. The novel cell lines and data from these experiments will facilitate developing treatments for CF individuals currently lacking an approved CFTR modulator therapy.

## Results

### Generation of novel Bmi-1/hTERT nasal and bronchial cell lines

Primary cell samples were transduced with a lentivirus containing mouse Bmi-1 and hTERT separated by the T2A self-cleaving peptide sequence (Figure 1A). By these methods, we generated two non-CF and three CF nasal cell lines, as well as two CF bronchial cell lines (summarized in Table I). Developing cell lines from primary tissue has been critiqued as inefficient because cells are lost during the selection of positive clones (19). Thus, to preserve patient material, an antibiotic selection cassette was not included in the vector design. Instead, transduced cells were gradually selected over time by the growth advantage conferred by Bmi-1/hTERT co-expression. Successful vector integration was confirmed at passage 6 (P6) and P15 by hTERT activity (Figure 1B) and Bmi-1 protein expression (Figure 1C-F). Notably, Bmi-1 expression was high in low passage primary cells but waned by P5. In a representative nasal and bronchial cell line (UNCNN2T and UNCCF9T, respectively), Bmi-1 expression was high at P15, confirming positive clonal selection. Mycoplasma-specific enzymes were undetectable in Bmi-1/hTERT cells between P6 and P7 or in the parent primary cells at P2. Short tandem repeat (STR) DNA profiling indicates each cell line is unique among human cell lines and samples present in Cellosaurus (v1.4.4). Further, the STR profile of each cell line matched that of the parent cells.

**Figure 1.**
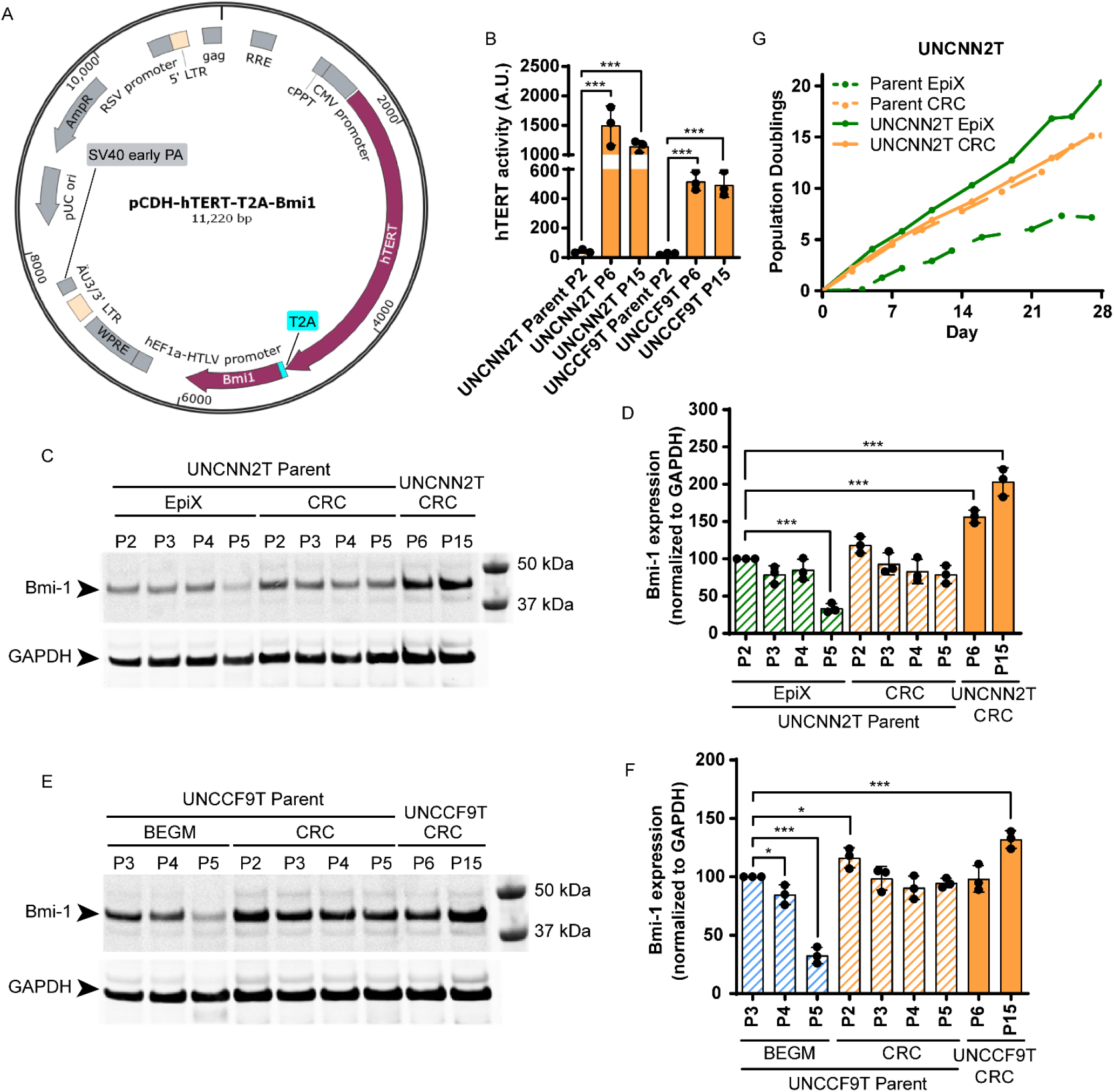
Generation of novel Bmi-1/hTERT nasal and bronchial cell lines. A) Vector map of the pCDH lentiviral plasmid containing mouse B cell-specific Moloney murine leukemia virus integration site 1 (Bmi-1) and human telomerase reverse transcriptase (hTERT) sequences linked by the T2A self-cleaving peptide sequence. B) Telomerase activity assay in the UNCNN2T nasal and UNCCF9T bronchial cell lines compared with parent cells. N=3 per condition. Log-transformed data were analyzed using an ordinary linear model with Tukey post test. C) Expression of Bmi-1 in the UNCNN2T nasal cell line and parent primary cells by western blot. D) Quantitation of (C); N=3. Data were analyzed using an ordinary linear model. E) Expression of Bmi-1 in the UNCCF9T bronchial cell line and parent primary cells by western blot. F) Quantitation of (E); N=3. Data were analyzed by using ordinary linear model. G) Growth curve data for the UNCNN2T cell line and parent primary cells in EpiX media and conditionally reprogrammed cell (CRC) growth conditions. All data presented as mean ± standard deviation (SD). * = p<0.05; *** = p<0.001.

**Table I.** Primary cell donor demographics. Cell line generated, cell source, *CFTR* genotype, and sex of two non-CF and five CF donors. Non-CF donors were not genotyped (indicated by a ‘-’’).

| Donor | Cell Line | Cell Source | *CFTR* Genotype | Sex |
| --- | --- | --- | --- | --- |
| Non-CF Donor 1 | UNCNN1T | Nasal curettage | - | Male |
| Non-CF Donor 2 | UNCNN2T | Nasal curettage | - | Female |
| CF Donor 1 | UNCX2T | Nasal curettage | W1282X/W1282X | Female |
| CF Donor 2 | UNCX3T | Nasal curettage | F508del/S492F | Male |
| CF Donor 3 | UNCX4T | Nasal curettage | F508del/F508del | Male |
| CF Donor 4 | UNCCF9T | Transplant lung | W1282X/W1282X | Female |
| CF Donor 5 | UNCCF10T | Transplant lung | W1282X/W1282X | Female |

### Bmi-1/hTERT nasal and bronchial cell lines exhibit enhanced growth properties

Traditionally, HNECs have been cultured using the same methods as HBECs (20), with expansion in bronchial epithelial growth medium (BEGM). Conditionally reprogrammed cell (CRC) technology improves HNEC growth capacity (21–23). However, the CRC technique requires co-culture with irradiated or mitomycin-treated NIH3T3 feeder cells. We found that nasal cells could also be expanded in EpiX media (Propagenix) as a feeder-free alternative to the CRC method (Figure 1G). To determine whether Bmi-1 and hTERT expression enhances HNEC growth properties, we compared the UNCNN2T nasal cell line and its parent cells in both CRC and EpiX culture conditions (Figure 1G). The Bmi-1/hTERT cell line vastly outgrew its parent nasal cells when cultured in EpiX media, whereas CRC culture promoted robust HNEC expansion regardless of Bmi-1/hTERT expression. The cell growth properties of all other nasal cell lines generated here can be found in the online supplements (Supplemental Figure 1A-D). Notably, the Bmi-1/hTERT bronchial cell lines could be effectively expanded using BEGM as previously described (15) (Supplemental Figure 1E-F). The cell morphology and electrophysiology of nasal cells cultured in EpiX or by CRC are compared in the online supplements (Supplemental Figure 1G-K).

### Pneumacult ALI promotes Bmi-1/hTERT cell line differentiation and ion transport function at passage 15

Nasal cell morphology and function is contingent upon *in vitro* culture conditions and media composition (24, 25). To optimize the differentiation of our Bmi-1/hTERT nasal cell model, we differentiated P15 UNCNN2T cells in UNC air-liquid interface (ALI) (26), Vertex ALI (27), and Pneumacult ALI (STEMCELL Technologies) media (Supplemental Figure 2A-E). The bioelectric properties were dramatically different between the three media conditions, with basal short-circuit current (Isc) averaging 7.1 ± 11.9 μA/cm^2^, 9.6 ± 1.6 μA/cm^2^, and 27.7 ± 7.9 μA/cm^2^ in UNC ALI, Vertex ALI, and Pneumacult ALI, respectively (mean ± SD). The apical addition of amiloride inhibited Isc by approximately 3%, 97%, and 29% in UNC ALI, Vertex ALI, and Pneumacult ALI, respectively. *In vivo* electrophysiology, as measured by nasal potential difference, is characterized by partial amiloride-sensitive Na+ transport (~50% basal Isc) (28). Thus, *in vivo* nasal electrophysiology is not perfectly modeled by any of the differentiation medias tested here. However, Pneumacult ALI is the closest match. Whole-mount immunostaining revealed a well-ciliated, pseudostratified epithelium with Pneumacult ALI whereas UNC ALI and Vertex ALI produced a thinner, poorly ciliated epithelium. Media comparisons with three published non-CF bronchial cell lines (15) yielded similar results (Supplemental Figure 3A-F). The ultimate goal of this study is to develop cell lines that can be used to evaluate CFTR rescue. Thus, we assayed CFTR correction by VX-445 and VX-661 (Elexacaftor/Tezacaftor; components of Trikafta) in three published bronchial cell lines with a F508del/F508del genotype (15) (Supplemental Figure 3G-J). The greatest CFTR response was again seen with Pneumacult ALI. Thus, Pneumacult ALI differentiation medium was used for the remainder of the study for all nasal and bronchial Bmi-1/hTERT cell lines unless otherwise indicated.

### Bmi-1/hTERT nasal cell lines model primary HNEC morphology and function

Hematoxylin and eosin (H&E) and Alcian blue-periodic acid-Schiff (AB-PAS) staining revealed a pseudostratified mucociliary epithelium in P6 and P15 UNCNN2T cells that was morphologically similar to parent HNECs at P2 (Figure 2A). These results were confirmed by whole-mount immunostaining, which illustrated the presence of both MUC5AC-producing goblet cells (green) and α-tubulin^+^ ciliated cells (white) (Figure 2B-D). From these data, we concluded that Bmi-1/hTERT nasal cell lines at mid- and late-passages produce a well-differentiated mucociliary epithelium that reflects primary cell morphology. Measurements with a 24-channel transepithelial current clamp amplifier (TECC-24) device demonstrated that parent HNEC electrophysiology was also recapitulated by the UNCNN2T cell line at mid- and late-passage, but with a significant increase in CFTR activity in P5 versus parent cells (Figure 2E-G). From this, we concluded that Bmi-1/hTERT nasal cell lines model primary HNEC morphology and function for at least 15 passages. Representative histology and whole-mount immunostaining of all other nasal cell lines are shown in the online supplement (Supplemental Figure 4).

**Figure 2.**
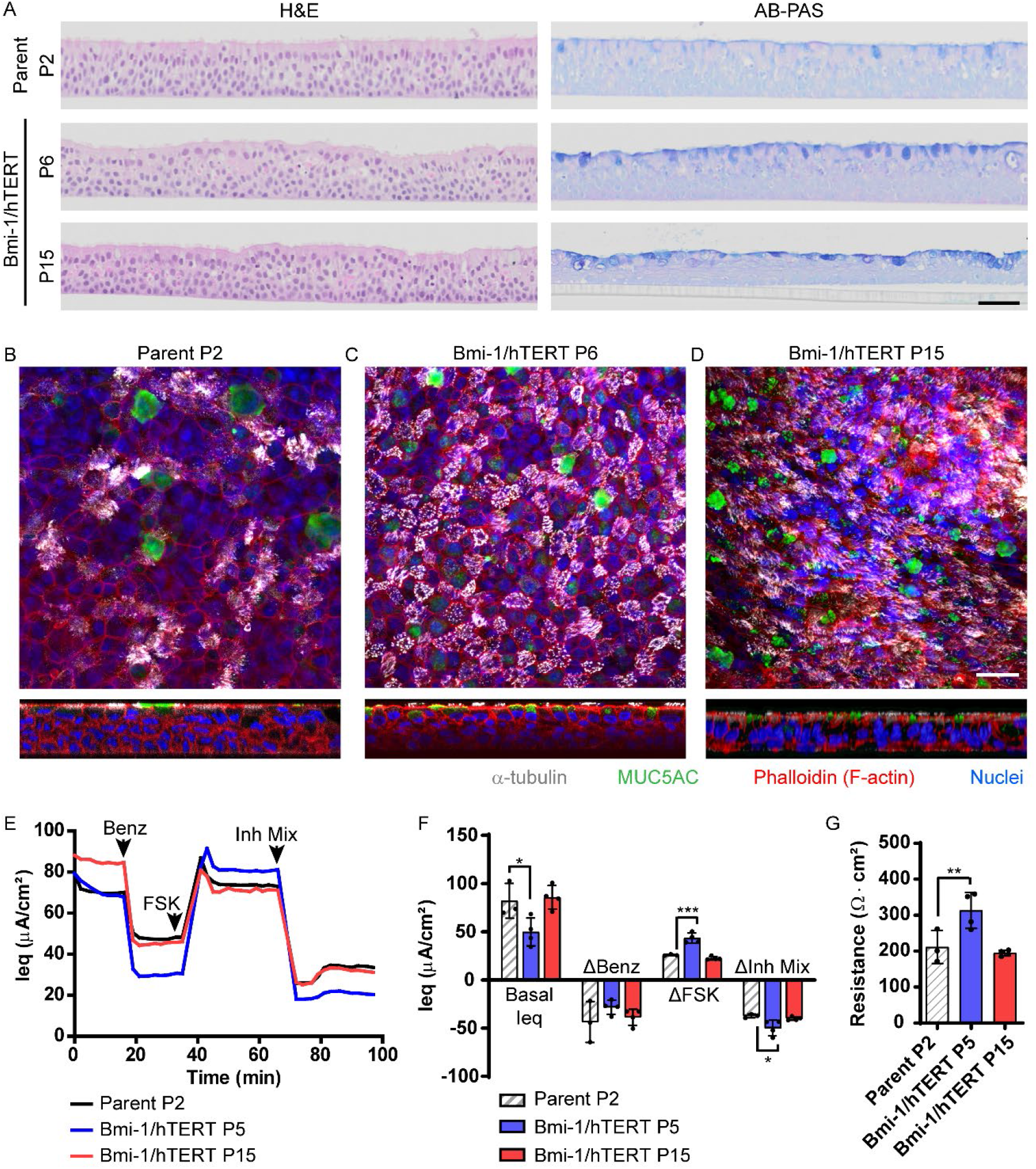
Nasal cell lines exhibit morphology and ion transport function representative of primary nasal cells. A) Hematoxylin and eosin (H&E) and Alcian blue-periodic acid-Schiff (AB-PAS) staining of UNCNN2T P2 parent cells and cell line at P6 and P15. Scale bar = 50 μm. B-D) Whole-mount immunostaining of UNCNN2T P2 parent cells (B), UNCNN2T P6 (C), and UNCNN2T P15 (D). α-tubulin (white), MUC5AC (green), Phalloidin (F-actin, red), and Hoechst (nuclei, blue). Scale bar = 25 μm. E-G) TECC-24 measurements of UNCNN2T P2 parent cells and cell line at P5 and P15. E) Representative TECC-24 tracing. F) Basal equivalent transepithelial current (Ieq) and change in Ieq (ΔIeq) in response to Benzamil (Benz), Forskolin (FSK), and an inhibitor mix (Inh Mix) consisting of CFTR inhibitor-172 (CFTRinh-172), GlyH-101, and Bumetanide. N=3-4. Data were analyzed using an ordinary linear model. G) Baseline resistance values. N=3-4. Log transformed data were analyzed using an ordinary linear model. All data presented as mean ± SD. * = p<0.05; ** = p<0.01; *** = p<0.001.

### Bmi-1/hTERT nasal cell lines predict the primary nasal cell response to CFTR modulators

The ultimate goal of developing patient-derived cell lines is to predict the clinical response to CFTR-targeted therapies. To test the ability of Bmi-1/hTERT nasal cell lines to predict CFTR modulator responses, we treated our F508del homozygous cell line, UNCX4T, with VX-445 and VX-661 (both at 5 μM). We also tested a recently described drug combination (29) which combines the CFTR corrector VX-809 with two novel small molecule correctors, 3151 and 4172 (each at 5 μM). Here, we call this treatment “triple corrector combination,” or “3C.” UNCX4T cells were treated with VX-445/VX-661, 3C, or a vehicle control (DMSO) and were assayed for CFTR function using a TECC-24 device (Figure 3A, C). VX-445/VX-661 and 3C rescued 23.0 ± 1.4% and 19.5 ± 4.8% of wildtype CFTR function, respectively (mean ± SD), determined by dividing the UNCX4T forskolin (FSK) response by the average response of non-CF nasal cell lines (46.5 ± 5.0 μA/cm^2^) (Figure 2F; Supplemental Figure 1H). These data align with preclinical studies of VX-445/VX-661 and clinical observations of Trikafta in F508del homozygous populations (30). VX-445/VX-661 and 3C treatment also significantly rescued CFTR ion transport in the paired parent nasal cells compared with a DMSO control (Figure 3B, C), rescuing 27.3 ± 5.1% and 26.7 ± 3.9% of wildtype CFTR function, respectively (mean ± SD) (Figure 3C). These data suggest that the novel CFTR corrector combination, 3C, might be as effective as Trikafta in rescuing F508del-CFTR and could represent a therapeutic candidate for those who cannot tolerate or do not respond to Trikafta treatment.

**Figure 3.**
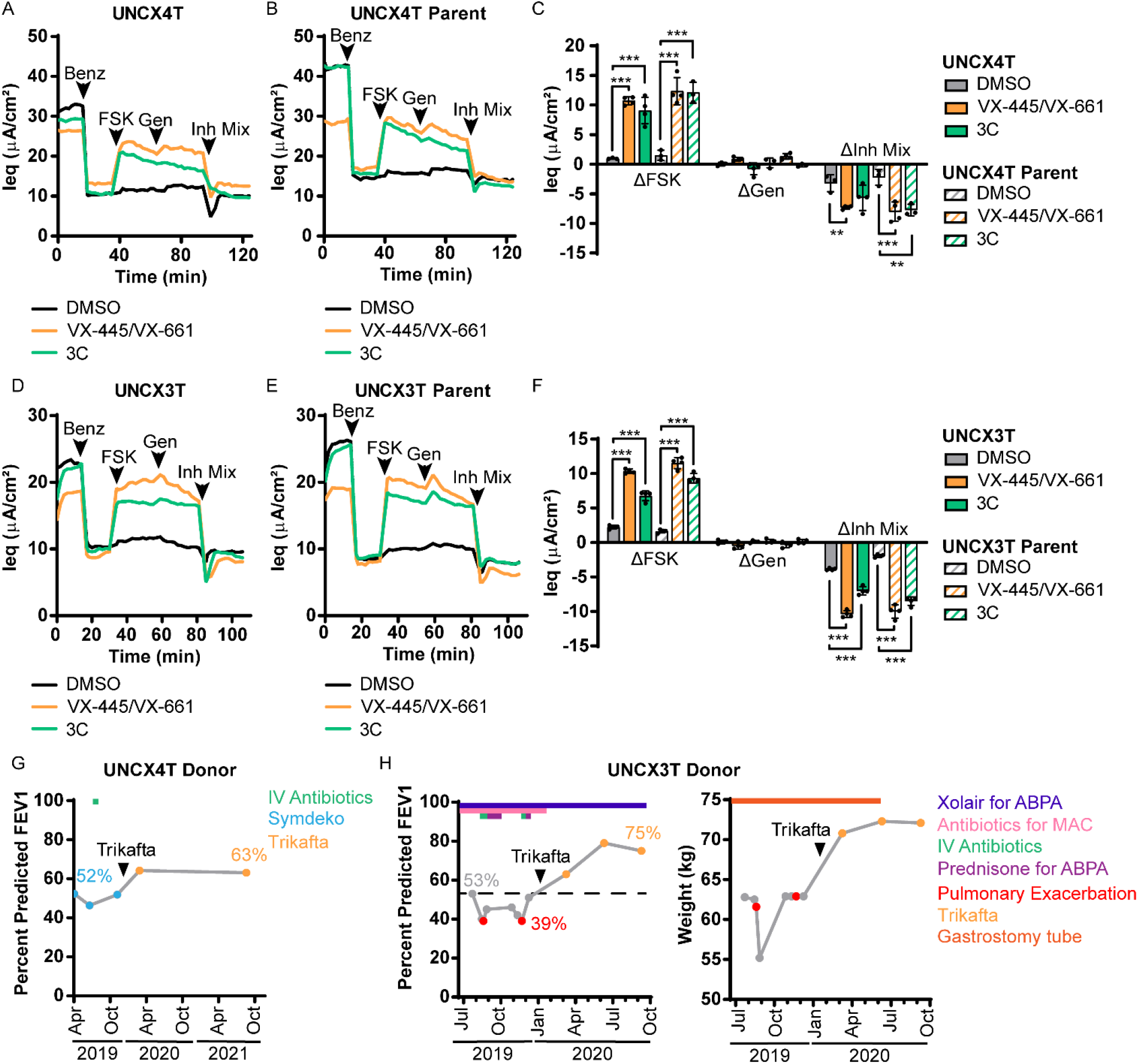
Nasal cell lines predict primary cell and clinical therapeutic response to CFTR modulators. A-B) Representative TECC-24 tracings of the F508del/F508del nasal cell line, UNCX4T (A), and parent primary cells (B) pretreated with DMSO, VX-445/VX-661, or a novel triple corrector combination (3C) for 48 hours. C) Δleq of UNCX4T and parent cells in response to FSK, Gen, and Inh Mix. N=3-4. Data were analyzed using an ordinary linear model. D-E) Representative TECC-24 tracings of the F508del/S492F nasal cell line, UNCX3T (D), and parent primary cells (E) pretreated with DMSO, VX-445/VX-661, or 3C for 48 hours. F) Δleq of UNCX3T and parent cells in response to FSK, Gen, and Inh Mix. N=3-4. Data were analyzed using an ordinary linear model. G-H) Change in the percent predicted forced expired volume in one second (FEV1) after Trikafta initiation in the UNCX4T donor (G) and the UNCX3T donor (H). Blue data points indicate FEV1 measured during Symdeko therapy, and orange data points indicate FEV1 measured during Trikafta therapy. FEV1 measured during a CF exacerbation is indicated by red data points. Treatment timeline for the UNCX4T and UNCX3T donors indicated above the FEV1 plot and includes timeline of intravenous (IV) antibiotics (green), Xolair for allergic bronchopulmonary aspergillosis (ABPA; dark blue), antibiotics for treatment of *Mycobacterium avium* complex (MAC; pink), and prednisone for ABPA (purple). I) Change in weight in kilograms (kg) in the UNCX3T cell donor after Trikafta initiation. Gastrostomy tube use and subsequent removal indicated by an orange bar. All data presented as mean ± SD. ** = p<0.01; *** = p<0.001.

Next, we assessed our F508del/S492F compound heterozygous cell line, UNCX3T, for its response to VX-445/VX-661, 3C, or a DMSO control (Figure 3D, F). We found that UNCX3T responded well to VX-445/VX-661 treatment, recapitulating 22.3 ± 0.7% of wildtype CFTR function (mean ± SD) (Figure 3F). However, 3C treatment was less effective, producing 14.6 ± 1.5% of wildtype function (mean ± SD). These findings were recapitulated in the UNCX3T parent cells (Figure 3E, F). Because 3C treatment produced less CFTR functional response in UNCX3T (one copy of F508del) than in UNCX4T (two copies of F508del), we posit that this modulator combination does not rescue S492F-CFTR as effectively as F508del-CFTR. Even so, the response to 3C falls well within the therapeutic window (i.e., 10% of wildtype CFTR function over baseline) and could serve as an alternative therapy. From these studies, we concluded that Bmi-1/hTERT nasal cell lines generated from CF donors accurately predict the primary cell response to FDA-approved and novel CFTR modulators.

### Nasal cell lines predict the clinical response to Trikafta in two patients with CF

All nasal curettage samples were obtained prior to the 2019 FDA approval of Trikafta. At the time of cell collection, the UNCX4T cell donor was prescribed tezacaftor/ivacaftor (trade name Symdeko), whereas the UNCX3T cell donor was not eligible for CFTR modulators. After nasal cell harvest and cell line generation, both donors became eligible for Trikafta and initiated treatment. Trikafta therapy was highly effective in these individuals, with a 11% and 22% increase in the percent predicted forced expired volume in one second (FEV1) over a six-month baseline in the UNCX4T and UNCX3T donors, respectively (Figure 3G-I). For the UNCX3T donor, Trikafta initiation correlated with other significant changes in health, including cessation of omalizumab (trade name Xolair) treatment for allergic bronchopulmonary aspergillosis (ABPA), reduced frequency of hospital admissions and intravenous (IV) antibiotics, and gastrostomy tube removal due to improved weight gain and retention (Figure 3H-I). Overall, the robust functional response to Trikafta that was observed in the UNCX4T and UNCX3T nasal cell lines *in vitro* was predictive of the cell donors’ positive clinical response to therapy.

### W1282X-CFTR is rescued by CC-90009, a novel cereblon modulator that promotes PTC readthrough

Current therapies are not effective in rescuing the truncated protein generated by the W1282X-CFTR variant. One proposed treatment strategy is to promote ribosomal readthrough of the PTC to generate full-length protein (31, 32). Low levels of readthrough can be accomplished *in vitro* with high concentrations of aminoglycosides (33). However, clinical readthrough agents are largely ineffective (34, 35) likely due to NMD, a surveillance pathway by which the cell detects and eliminates PTC-containing mRNA transcripts (36). Indeed, a recent study found substantial degradation of the CFTR transcript in an individual homozygous for the W1282X variant, with the mutated CFTR expressed at only 2.1% of wildtype levels (37). Thus, many groups have hypothesized that effective treatment of nonsense mutations will also require NMD inhibition (31, 37, 38). Studies that bypass NMD by expressing intron-less complementary DNA (cDNA) copies of W1282X-CFTR have demonstrated that the truncated protein exhibits defective cellular trafficking and gating which can be augmented by CFTR modulators (39, 40). Yet in primary cells, CFTR modulators alone do not rescue function (32). We therefore hypothesized that effective rescue of W1282X-CFTR would require a combination of therapeutic approaches to: 1) promote PTC readthrough, 2) inhibit NMD, and 3) modulate the resulting CFTR protein.

Recently, a novel class of cereblon (CRBN) E3 ligase modulators have been shown to significantly improve PTC readthrough by aminoglycoside compounds (18). One of these agents, CC-90009, is currently under investigation for treatment of relapsed and refractory acute myeloid leukemia (AML) in a phase 1 clinical trial (NCT02848001). CC-90009 mediates the interaction between CRBN and the G1 to S phase transition 1 (GSPT1) protein (also known as the eukaryotic release factor 3a; eRF3a), which functions as a key player in stop codon recognition and translation termination. Upon interaction with CRBN, GSPT1 is ubiquitinated and targeted for proteasomal degradation. Thus, we hypothesized that CC-90009 would further enhance therapeutic readthrough and rescue of W1282X-CFTR.

To test this, we combined an inhibitor of NMD (0.3 μM Smg1i), an aminoglycoside (10 μM G418), and a CFTR corrector (3 μM VX-809) with and without 10 μM CC-90009 and compared both combinations to a vehicle control (DMSO) in the W1282X/W1282X nasal cell line, UNCX2T (Figure 4A-B). CFTR function (i.e., FSK response) was undetectable in vehicle-treated cells, whereas treatment with Smg1i, G418, and VX-809 partially restored channel function to 3.4 ± 1.6% of wildtype levels (mean ± SD). CFTR function was significantly increased by the inclusion of CC-90009 to 27.0 ± 10.5% of wildtype (mean ± SD). To understand this dramatic increase, we studied each compound individually and found that 10 μM CC-90009 alone effectively rescued CFTR function (Supplemental Figure 5A-C; Figure 4A-B). None of the other compounds tested (i.e., Smg1i, G418, or VX-809) provided significant functional rescue. CFTR mRNA transcript levels increased significantly after treatment with CC-90009, G418 (10 μM or 200 μM), or Smg1i as previously reported (31, 38) (Supplemental Figure 5D).

**Figure 4.**
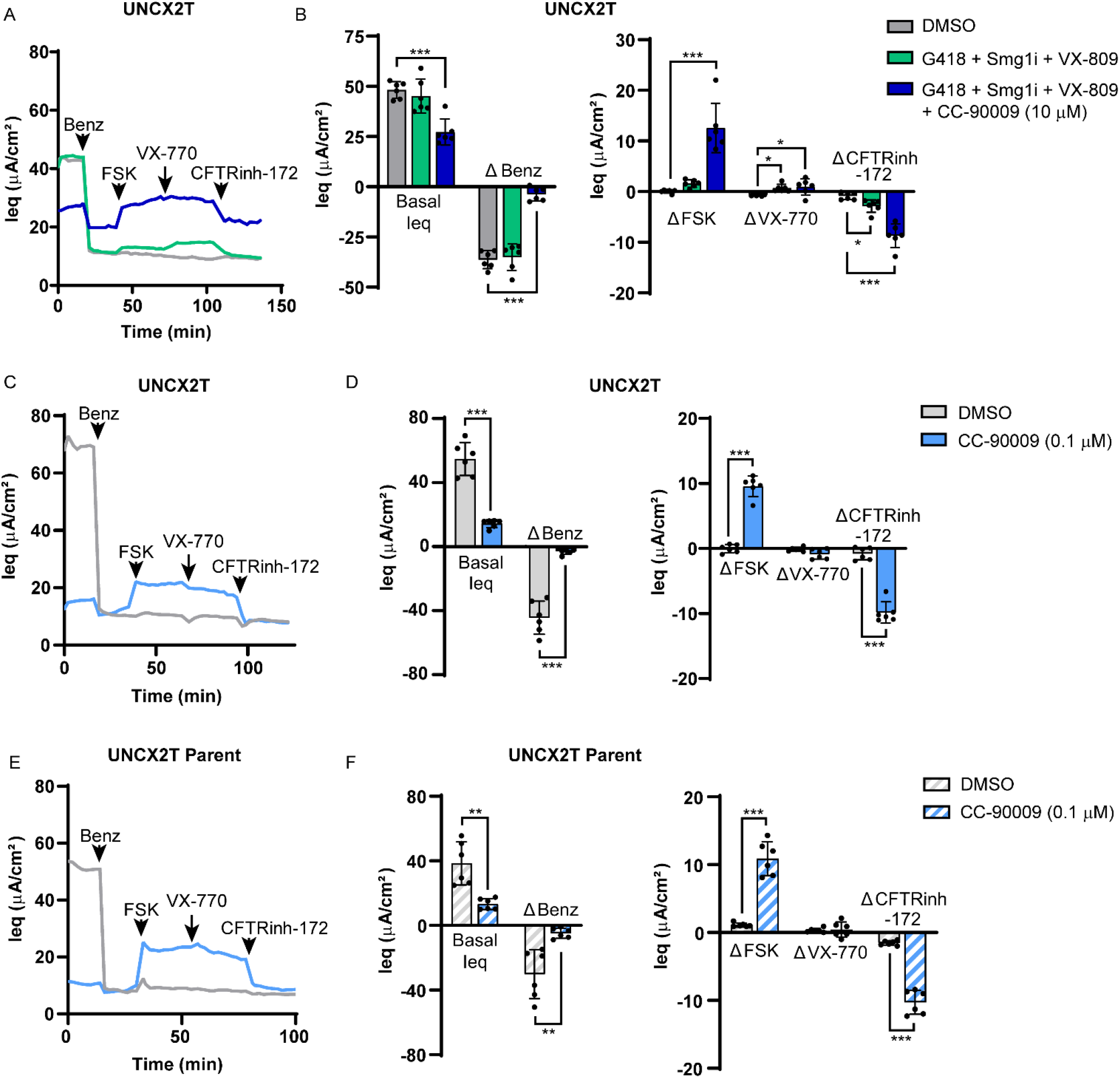
The G1 to S phase transition protein 1 (GSPT1) degrader CC-90009 rescues W1282X CFTR in a nasal cell line and parent primary cells. A) Representative TECC-24 tracing of the W1282X/W1282X nasal cell line, UNCX2T, pretreated with 10 μM G418 + 0.3 μM Smg1i + 3 μM VX-809 with and without 10 μM CC-90009 or DMSO for 48 hours. B) Basal Ieq and ΔIeq in response to Benz (left). ΔIeq in response to FSK, VX-770, and CFTRinh-172 (right). N=6. Data were analyzed using an ordinary linear model. C) Representative TECC-24 tracing of UNCX2T pretreated with 0.1 μM CC-90009 or DMSO for 24 hours. D) Basal Ieq and ΔIeq in response to Benz (left). ΔIeq in response to FSK, VX-770, and CFTRinh-172 (right). N=6. Data were analyzed using an ordinary linear model. E) Representative TECC-24 tracing of UNCX2T parent primary cells pretreated with 0.1 μM CC-90009 or DMSO for 24 hours. F) Basal Ieq and ΔIeq in response to Benz (left). ΔIeq in response to FSK, VX-770, and CFTRinh-172 (right). N=6. Data were analyzed using an ordinary linear model. All data presented as mean ± SD. * = p<0.05; ** = p<0.01; *** = p<0.001.

CC-90009 has been previously studied at 10 μM concentrations and was reported to have negligible cytotoxicity (18). Yet in our culture system, the transepithelial resistance was reduced by ~50% in cultures treated with 10 μM CC-90009 (Supplemental Figure 5C). To optimize dosing for airway epithelial cells, we treated primary nasal cells with escalating concentrations of CC-90009 from 0.1 to 1 μM. The lowest dose tested (0.1 μM) did not attenuate transepithelial resistance (Supplemental Figure 5E) or trigger cytotoxicity as measured by lactate dehydrogenase (LDH) release (Supplemental Figure 5F). Cell morphology was also not altered by 0.1 μM CC-90009 treatment (Supplemental Figure 5G). We then assessed GSPT1 protein levels by western blot and found substantial degradation at 0.1 μM (Supplemental Figure 5H-I).

Having established a safe and effective dose, we assessed the effect of 0.1 μM CC-90009 treatment on UNCX2T electrophysiology. At 0.1 μM, CC-90009 rescued approximately 20.6 ± 3.4% of wildtype CFTR function in the UNCX2T cell line (mean ± SD) (Figure 4C-D). These findings were validated in the UNCX2T parent nasal cells where 0.1 μM CC-90009 rescued 24.1 ± 5.5% of wildtype function (mean ± SD) (Figure 4E-F). We next tested CC-90009’s ability to rescue CFTR in the two W1282X/W1282X bronchial cell lines, UNCCF9T and UNCCF10T, and in the parent primary cells. CC-90009 rescued 22.8 ± 5.8% and 17.9 ± 5.6% wildtype CFTR function in the bronchial cell lines and primary cells, respectively (mean ± SD) (Figure 5A-D). Based on the literature, we suspect that the primary mechanism for W1282X-CFTR rescue is enhanced ribosomal readthrough (18). CFTR protein level is low in primary airway epithelial cells and made even lower by NMD in cells carrying PTC variants, making it challenging to assess. Importantly, we did not observe CC-90009 to increase CFTR activity in F508del- or wildtype-CFTR primary HBECs; rather, it was slightly reduced (Supplemental Figure 6). This supports the notion that CC-90009 specifically acts to promote PTC readthrough rather than by non-specific effects on cellular ion transport. We measured CFTR mRNA by qRT-PCR and found a significant increase in response to CC-90009 treatment (Figure 5E). This finding is consistent with the published observation that GSPT1 degradation stabilizes PTC-containing transcripts through a partial suppression of NMD (18). Overall, CC-90009 treatment significantly rescued CFTR function in a cohort of three W1282X homozygous patient-derived cell lines and primary nasal and bronchial epithelial cells.

**Figure 5.**
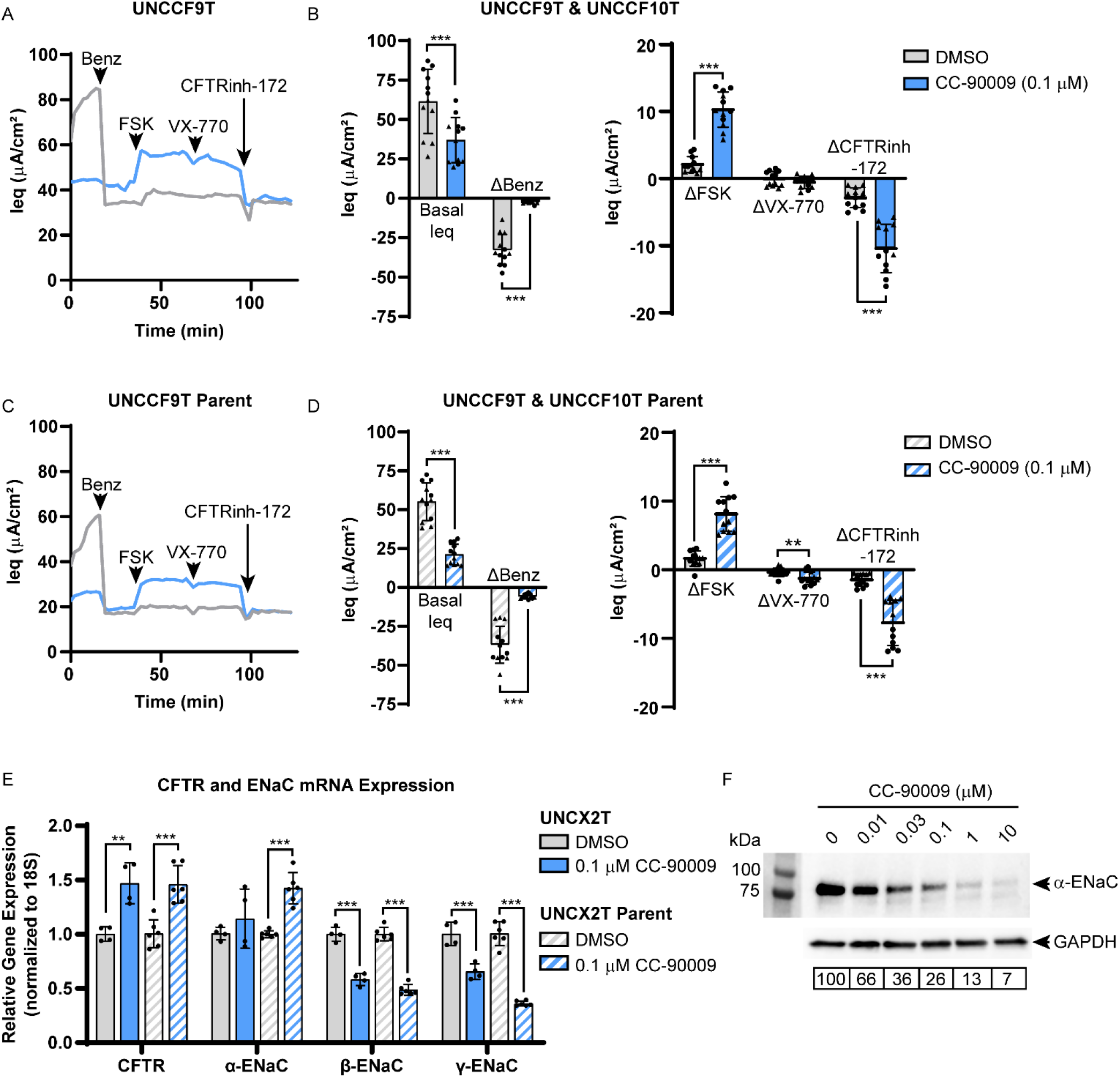
CC-90009 stabilizes W1282X CFTR mRNA and inhibits epithelial sodium channel (ENaC) activity in bronchial cell lines. A) Representative TECC-24 tracing of the W1282X/W1282X bronchial cell line, UNCCF9T, pretreated with 0.1 μM CC-90009 or DMSO for 24 hours. B) Basal Ieq and Δleq in response to Benz (left). Δleq in response to FSK, VX-770, and CFTRinh-172 (right) in the UNCCF9T (circles) and UNCCF10T (triangles) cell lines. N=6 for each cell line. Data were analyzed using a linear mixed-effects model with the donor as a random effect factor. C) Representative TECC-24 tracing of the UNCCF9T parent primary cells pretreated with 0.1 μM CC-90009 or DMSO for 24 hours. D) Basal Ieq and Δleq in response to Benzamil (left). Δleq in response to FSk, VX-770, and CFTRinh-172 (right) in the UNCCF9T (circles) and UNCCF10T (triangles) parent primary cells. N=6 for each donor. Data were analyzed using a linear mixed-effects model with the donor as a random effect factor. E) Relative CFTR and ENaC mRNA levels by qRT-PCR in UNCX2T and parent cells. N=4-6. Data were analyzed using an ordinary linear model. F) Western blot for α-ENaC in UNCCF9T parent cells pretreated with escalating doses of CC-90009 for 24 hours. Protein expression normalized to GAPDH and relative to the DMSO control quantified below. Cells were grown in Vertex ALI media to increase the level of ENaC expression to be detectable by western blot. All data presented as mean ± SD. ** = p<0.01; *** = p<0.001.

### CC-90009 diminishes epithelial sodium channel (ENaC) activity

ENaC is a heterotrimeric ion channel comprising three homologous subunits, α, β, and γ (41) and is responsible for the tight regulation of sodium and fluid absorption across epithelia (i.e., the lungs, colon, and kidneys). ENaC functionality and localization are regulated by several factors including hormones and proteases. Though the biology of ENaC is complex, ubiquitin ligases and proteasomal pathways play a known role (42).

We noted a striking and unexpected decrease in ENaC activity after treatment with CC-90009. The response to benzamil, a potent inhibitor of ENaC, was reduced by ~80-95% across all cell lines and parent cells treated with CC-90009 (Figure 4B, D, F; Figure 5B, D; Supplemental Figure 5B; Supplemental Figure 6B, D). The basal equivalent current (Ieq) was also reduced in treated cells, indicating reduced sodium absorption at baseline. GSPT1 is the only reported target of CC-90009 (43). Thus, ENaC or a regulator of ENaC represents a previously undescribed off-target effect of the compound. Treatment with CC-90009 decreased β- and γ-ENaC mRNA expression in both the primary nasal cells and nasal cell line while the effect on α-ENaC mRNA was cell-line dependent (Figure 5E). By western blot, we observed a dose-dependent decrease in α-ENaC protein levels to about one-third of baseline expression at 0.1 μM CC-90009 (Figure 5F). Thus, CC-90009 modulates ENaC expression at both the mRNA and protein levels. Though the effects of CC-90009 on ENaC were unexpected, these too may prove therapeutic. Increasing airway hydration through ENaC inhibition has been proposed previously as a treatment strategy for CF (44). Future mechanistic work will be necessary to discern the link between CC-90009 and ENaC and may uncover novel pathways for ENaC ubiquitination.

In summary, we have shown that treatment with CC-90009 increases CFTR function and decreases ENaC expression in W1282X patient-derived cell lines and primary airway epithelial cells. GSPT1 degradation by CC-90009 represents a new therapeutic strategy for treating nonsense mutations in the CFTR transcript.

## Discussion

The extension of CFTR modulators to 90% of people with CF represents a tremendous victory. However, developing targeted therapies for the remaining 10% will be challenging. This challenge is heightened by the limited availability of rare-genotype primary cells. For this reason, the field has turned to cell lines for drug discovery efforts.

Preclinical studies rely heavily on Fischer rat thyroid (FRT) cells engineered to express high levels of mutant CFTR (9, 45, 46). While highly informative, FRT cells do not capture the spectrum of response to CFTR modulators that is routinely observed in clinical trials, even in patient cohorts with the same genotype (6, 7, 47, 48). The inability of FRT cells to mimic patient-specific differences in CFTR regulation and function is an inherent shortcoming of the model (1, 27). Further, because of its robust CFTR overexpression, the FRT model has a greater chance of producing false-positive “hits” during drug screens (19). This was documented in a study of the W1282X variant in which the modulators VX-809 and VX-770 rescued CFTR function in the FRT model but not in patient-derived nasal epithelial cells (39). FRT and other overexpression cell lines may not accurately recapitulate mutations that cause a pathologic decrease in CFTR transcript level.

Though primary HBECs are the gold standard of CF disease modeling, they can only be expanded for ~10 population doublings using conventional growth methods (10) or ~25 doublings using newer CRC culture methods (12, 49). HNECs also exhibit extended growth capacity using CRC technology (21–23). However, with extensive population doublings comes squamous transformation and a reduction in ion transport (12). Here, we describe the development of Bmi-1/hTERT cell lines derived from primary airway epithelial cells that can be expanded for at least 30 population doublings (15 passages) while recapitulating primary cell morphology and CFTR function. The ability to amply expand and assay CFTR function in patient-derived cells represents a major advancement.

In this study, we directly compare the functional response of patient-derived Bmi-1/hTERT cell lines and the primary cells from which they were developed. Across five CF donors (three nasal and two bronchial) carrying a range of *CFTR* mutations and using both FDA-approved and novel CFTR modulators, we found that patient-derived cell lines accurately predicted primary cell CFTR rescue. We created two nasal epithelial cell lines from donors carrying the F508del allele. These cell lines served as a positive control for effective modulator response (i.e., Trikafta) and provided proof-of-concept that patient-derived cell lines are predictive of clinical response. One of the two F508del-carrying cell lines, UNCX3T, was created from a compound heterozygous donor with genotype F508del/S492F. S492F encodes a rare mutation in the nucleotide-binding domain 1 (NBD1) of *CFTR* (9), with only 24 instances registered on CFTR2.org. The UNCX3T cell line provides proof-of-concept that rare *CFTR* variants can also be studied in the compound heterozygous context. Notably, the ongoing global coronavirus disease 2019 (COVID-19) pandemic began shortly after FDA approval of Trikafta in November 2019 and dramatically changed the way that CF clinic visits are conducted. As such, the UNCX4T donor has only two post-Trikafta measurements recorded. Even so, this small N=2 patient cohort demonstrates that the response of Bmi-1/hTERT nasal cell lines aligns with clinical measurements.

Nonsense mutations in the *CFTR* transcript produce little to no CFTR protein. Thus, therapeutic efforts have logically focused on promoting ribosomal readthrough. A small study demonstrated that short-term intranasal administration of the aminoglycoside, gentamicin, partially reduced nasal potential difference in CF patients harboring nonsense mutations (50, 51). However, the known retinal toxicity, ototoxicity, and nephrotoxicity of aminoglycosides hinders clinical translation (52–55). Ataluren (PTC124), a non-aminoglycoside inducer of readthrough, was recently assessed in a phase 3 clinical trial for treatment of CF nonsense mutations. However, the primary and secondary endpoints of change in FEV1 and rate of pulmonary exacerbations were not met, and no further clinical development for CF is planned (34).

Translation termination begins when the eukaryotic release factor 1 (eRF1) binds the ribosomal decoding A site and recognizes a stop codon (56). Through its interaction with GSPT1, eRF1 hydrolyzes the peptidyl-tRNA and mediates the release of nascent polypeptide (57). *In vitro* reporter assays have demonstrated that depletion of eRF1 or GSPT1 by short interfering RNAs (siRNAs) or antisense oligonucleotides (ASOs) promotes PTC readthrough (58, 59). This work was confirmed by Huang et al., who assessed PTC readthrough in a murine hemophilia model by eRF1 and GSPT1 knockdown with ASOs (60). Only a modest increase in PTC readthrough was observed. However, the group reported striking levels of synergy with aminoglycoside treatment. Recently, a study by Sharma et al., demonstrated PTC readthrough in CFTR after small molecule knockdown of eRF1 (61). This group reported aminoglycoside synergy in FRT and 16HBE gene-edited cell models. However, when tested in primary bronchial cells, the group observed only low levels of correction. In this study, we explore a novel class of compounds that degrades the translation termination factor GSPT1.

A new class of small molecules targeting GSPT1 has been recently described as potent tumoricidal agents, with activity against AML (62–67). Baradaran-Heravi et al. was the first group to assess these small molecule GSPT1 degraders for ribosomal readthrough and rescue of nonsense mutations (18). In a panel of human disease models (mucopolysaccharidosis type I-Hurler, late infantile neuronal ceroid lipofuscinosis, Duchenne muscular dystrophy and junctional epidermolysis bullosa), the authors found that CC-90009 and analog CC-885 act synergistically with aminoglycosides to promote PTC readthrough (18). This was confirmed by evaluating readthrough after siRNA knockdown of GSPT1 (18). The similarity in effect between CC-90009 treatment and GSPT1 siRNA provides strong evidence that observed PTC readthrough can be attributed to reduced GSPT1 levels rather than an off-target effect of the compound. This was confirmed by Surka and colleagues who demonstrated that GSPT1 degradation is both necessary and sufficient to account for CC-90009’s activity in AML cell lines (43). Similarly, we found that CC-90009 treatment produced a robust dose-dependent decrease in GSPT1 protein in patient-derived airway epithelial cells.

Unlike Huang et al. and Baradaran-Heravi et al., we observed high levels of PTC readthrough by GSPT1 degradation alone with no apparent aminoglycoside synergy. This difference could be due to cell-type specific differences in GSPT1 expression or the unique biology of CFTR. We previously reported a robust non-linear relationship between CFTR function and the proportion of CFTR-expressing cells (68). Accordingly, even low levels of PTC readthrough could account for the ~20% of wildtype CFTR function recovered by CC-90009 treatment.

A weakness of this study is the inability to detect CFTR protein expression in PTC-carrying primary airway epithelial cells. Even in non-CF cells, CFTR protein abundance is low and CFTR is barely detectable after immunoprecipitation with a CFTR antibody. Studies in progress are evaluating more sensitive methods to detect changes in CFTR protein expression. Despite current CFTR protein evidence, the lack of therapeutic effect on F508del- and wildtype-CFTR (Supplemental Figure 6) strongly suggests that CC-90009’s activity is specific to PTC-relevant pathways.

Partial suppression of NMD by ribosomal readthrough with aminoglycosides (69, 70) or with CC-90009 (18) is a well-established phenomenon. Likewise, we found that CC-90009 increased the level of CFTR mRNA expression (Figure 5E). However, a far greater increase was seen after treatment with Smg1i (Supplemental Figure 5D) with no corresponding increase in CFTR ion transport (Supplemental Figure 5A-B). Thus, we posit that the functional rescue of W1282X-CFTR is likely not explained by NMD suppression alone. Follow-up proteomic studies are required to definitively confirm PTC readthrough as the primary mechanism of W1282X-CFTR rescue.

CC-90009 reportedly has little effect on the proteome apart from GSPT1 (64, 67). However, Aliouat et al., documented over 2000 genes with significant changes in mRNA expression and protein translation after GSPT1 knockdown (71). In the present study, we report an unexpected decrease in ENaC function as a result of CC-90009 treatment. These findings could have implications for the ongoing clinical evaluation of CC-90009 for treatment of AML (NCT02848001). Early clinical trial results reported hypotension as a dose-limiting toxicity and a treatment-emergent adverse event (65). Sodium reabsorption via ENaC is known to determine extracellular fluid volume and regulate blood pressure (72, 73). Indeed, human disease caused by ENaC gain-of-function or loss-of-function in Liddle syndrome or Pseudohypoaldosteronism type I (PHA-1), respectively, provide strong evidence for a direct link between ENaC function and systemic blood pressure, with ENaC overactivity leading to hypertension and reduced activity leading to hypotension (74–76). Thus, we posit that the adverse hypotension observed in the CC-90009 clinical trials could be explained by off-target ENaC inhibition. We speculate that the decreased level of β- and γ-ENaC may destabilize the heterotrimer and cause the loss of α-ENaC protein. Future work to understand the connection between CC-90009 and ENaC will be necessary.

One concern of therapeutically degrading GSPT1 is the possibility of normal termination codon (NTC) readthrough. However, NTCs are thought to be more highly regulated than PTCs since basal levels of readthrough are ~10 times more likely at a PTC (60). Further, studies of aminoglycoside- and Ataluren-induced readthrough demonstrate no adverse effect on translation termination at NTCs (53, 77). While these studies set the precedent that readthrough therapies are safe, future work will be required to determine whether GSPT1 degradation affects translation termination at NTCs.

In this study, we describe a pipeline for developing patient-derived airway epithelial cell lines to model rare CFTR variants. Bmi-1/hTERT nasal and bronchial cell lines are highly representative of primary airway cells and predictive of clinical response to CFTR modulators. We present the novel finding that GSPT1 degradation by CC-90009 promotes robust functional rescue of W1282X-CFTR. Follow-up studies to understand the mechanism of rescue will be vital to develop this new therapeutic strategy for treating people with CF currently ineligible for modulator therapy.

## Methods

### Primary cell isolation and tissue culture

Primary HNECs were obtained by nasal curettage from two non-CF and three CF donors, yielding 2.1 × 10^6^ cells on average. Demographics and CFTR genotypes are summarized in Table I. Freshly obtained samples were transferred into Lactated Ringer’s Solution. Tissue dissociation was performed by treating with dithiothreitol (DTT, 0.5 mg/mL; Sigma-Aldrich, D0632) and DNase (10 μg/mL; Sigma-Aldrich, DN25) for 15 minutes. Cells were disaggregated by incubation with Accutase (Sigma-Aldrich, A6964) for 10 minutes. HBECs from explanted CF transplant lungs were obtained and isolated as previously described (26). HNECs and HBECs were cultured by CRC culture methods (12, 49) unless otherwise indicated. Briefly, cells were co-cultured with irradiated NIH3T3 fibroblasts in the presence of the rho-kinase inhibitor, Y-27632. Primary cells were cultured with an antibiotic/antifungal cocktail for the first two days.

### Production and titering of the Bmi-1/hTERT lentivirus

HEK 293T cells were thawed and passed a minimum of two times in p100 dishes with Dulbecco’s Modified Eagle’s Medium (DMEM) + 10% Fetal Bovine Serum (FBS) + 1% Penicillin/Streptomycin before transfecting the cells at 85-95% confluence with 8 ng pCDH-hTERT-T2A-Bmi-1, 8 ng psPAX, and 4 ng VSVG plasmids, using Lipofectamine 2000 (Invitrogen, 11668-019) and following the manufacturer’s protocol. Cells were maintained at 37°C and 5% CO_2_. Media was changed and collected every 24 hours for up to 72 hours and stored at 4°C. A portion of virus-containing medium was aliquoted and frozen at −80°C, and the remainder was concentrated with the Lenti-X Concentrator kit (Clontech, 631231). Viral titering was performed using 15 μL of unconcentrated virus or 1.5 μl of concentrated virus with the Lenti-X qRT-PCR Titration Kit (Clontech, 632165). Results were read on an ABI QuantStudio 6 RT-PCR machine.

### Creation of Bmi-1/hTERT growth-enhanced cell lines

Primary cells were cultured in CRC conditions to P1. At ~30% confluence, irradiated NIH3T3 cells were removed with trypsin and the media was replaced with irradiated NIH3T3-conditioned media (78) containing polybrene (10 μg/mL) and hTERT-T2A-Bmi-1 lentivirus (Figure 1A) at a multiplicity of infection (MOI) of 2. After 24 hours, cells were washed with 1X PBS and transduced again with fresh virus to increase transduction efficiency. After 48 hours of lentiviral exposure, CRC culture conditions were restored by adding irradiated NIH3T3 cells. Cells were passed using Accutase at 70-90% confluence. Growth curves were created by calculating population doublings as the log base 2 of the ending cell number divided by the starting cell number.

### Validation of hTERT expression and activity

hTERT expression was determined using quantitative analyses of telomerase activity by the combination of the conventional telomeric repeat amplification protocol (TRAP) with real-time quantitative PCR (RT-qPCR) (79, 80). Cells were lysed with a buffer containing 0.5% CHAPS [3-((3-chloamidopropyl)-dimethyl-ammonio)-1-propanesulonate], 10 mM Tris-HCl, pH7.5, 1 mM MgCl_2_, 1 mM EGTA, 0.1 mM Benzamidine, 10 U/ml RNasin and 10% glycerol. Protein concentration was measured by DC protein assay (BioRad, 500-0112), and 5 mM β-mercaptoethanol was added to the samples. Each 40 μl reaction contained 2 μg of cell lysate diluted in 0.1 mg/ml BSA, 1X TRAP reaction buffer [20 mM Tris-HCl (pH 8.3), 1.5 mM MgCl_2_, 10 mM EGTA, and 50 mM KCl, 50 μM of each deoxynucleotide triphosphate, 910 nM TS forward primer (see sequence below), and 0.4 μg of T4 gene protein (NEB, M0300S)]. The reaction mixture was incubated at 33°C for 5 hours followed by 95°C for 10 minutes to inactivate telomerase. RT-qPCR with iQ SYBR Green Supermix (Bio-Rad,170-8882) was then used to quantitate the number of substrate molecules to which telomeric repeats were added. Each 25 μl reaction contained 300 nM TS forward and reverse primer and 1 μl of the product from the first step. A standard curve was produced using 10X serial dilution of HEK293 cell extracts, from 4 μg to 4 ng. All samples were run in triplicate. The sequences of the TS primers are 5’-AATCCGTCGAGCAGAGTT (forward) and 5’-CCCTTACCCTTACCCTTACCCTAA (reverse).

### Western blot analysis

Cells were rinsed with cold PBS and lysed in RIPA buffer (Boston Bioproducts, BP-115D) with a Halt protease and phosphatase inhibitor cocktail (Thermo Fisher Scientific, 1861280) for 30 minutes on ice and spun at 16,000 g for 15 minutes to remove the insoluble fraction. Protein concentration was determined using the DC protein assay (Bio-Rad, 5000112). For each sample, 30 μg protein was loaded onto an SDS-PAGE gel and transferred to a nitrocellulose membrane. To validate Bmi-1 expression, blots were incubated with anti-Bmi-1 antibody (Cell Signaling, 5856S), followed by goat-anti-rabbit IgG (H+L) DyLight800 (Invitrogen, SA5-10036). Signals were detected and quantified by an Odyssey Image scanner (LI-COR). To determine the effect of CC-90009 on α-ENaC and GSPT1 expression, blots were incubated with anti-α-ENaC (mAb UNC1 19.2.1) and anti-GSPT1 antibody (Abcam, ab49878), followed by peroxidase-linked donkey anti-mouse IgG (Thermo Fisher Scientific, 45000679) and donkey anti-rabbit IgG (Thermo Fisher Scientific, 45000682), respectively. Signals were detected using Clarity Western ECL substrate (BioRad, 170-5061), imaged with Azure Image System (Azure Biosystems), and quantified with ImageQuant TL. To normalize protein loading, blots were re-probed with anti-GAPDH (Proteintech, 60004-1-Ig).

### RNA purification and RT-qPCR

Total RNA was purified using Direct-zol RNA mini-prep kit (Zymo Research, R2052) and quantified using a NanoDrop-8000 spectrophotometer (Thermo Fisher Scientific). Taqman assays for CFTR (Integrated DNA Tech., Hs.PT.58.28207352), α-ENaC/SCNN1A (Hs01013028_m1), β-ENaC/SCNN1B (Hs00165722_m1) and γ-ENaC/SCNN1G (Hs00168918_m1) were performed with the Luna Universal Probe One-Step RT-qPCR Kit (NEB, E3006L). Expression of each gene was normalized to 18S (Hs99999901_s1). All Taqman probes were purchased from Applied Biosystems except the CFTR probe. RT-qPCR was run on Applied Biosystem QuantStudio 6 (Thermo Fisher Scientific).

### Mycoplasma testing, DNA fingerprinting, and CFTR genotyping

Mycoplasma contamination was evaluated by the MycoAlert Plus Mycoplasma Detection Kit (Lonza LT07-710) with the MycoAlert Assay Control Set (Lonza LT07-518) according to the manufacturer’s instructions. Luminometry results were read on a FLUOstar Omega microplate reader (BMG LABTECH). Cell line authentication was performed by Labcorp’s Cell Line Testing Division on July 13, 2021, using the PowerPlex^®^ 16HS (Promega Corporation) for STR DNA profiling. *CFTR* genotyping was performed by clinically approved laboratories and retrieved from the clinical record.

### Culture at air-liquid interface

Cell culture inserts were coated with human placental type IV collagen (Sigma-Aldrich, C7521). CRC expanded cells were seeded at 2.5 × 10^5^ cells on 12-mm Snapwells (Corning Costar, 3407) or at 3.3 × 10^4^ cells on 6.5-mm Transwells^®^ (Corning Costar, 3470) or HTS Transwell^®^-24-well Permeable Support (Corning Costar, 3378) and transitioned to air-liquid interface (ALI) when cells were confluent. Cells were maintained in Pneumacult ALI medium (STEMCELL Technologies, no. 05001) for 21-32 days after seeding. For drug treatment, compounds were added in basolateral media, and 10 μl or 50 μl were added apically for 6.5 mm and 12 mm transwells, respectively.

### Whole-mount immunostaining and imaging

ALI cultures were fixed for 15 minutes on day 21-32 in 4% paraformaldehyde and stained for α-tubulin (Sigma-Aldrich, MAB1864; 3 μg/mL), MUC5AC (Thermo Fisher Scientific, 45M1; 4μg/mL), F-actin (Phalloidin; Invitrogen, A22287), and DNA (Hoechst 33342; Invitrogen, H3570) using species-specific secondary antibodies as previously described (81). Cultures were imaged using a Leica TCS SP8 with a 63X oil objective.

### Electrophysiology measurements

To select a nasal cell differentiation media, Ussing chamber analysis was performed with the addition of 100 μM amiloride (Amil), 10 μM forskolin (FSK), 10 μM CFTRinhibitor-172 (CFTRinh-172), and 10 μM uridine-5’-triphosphate (UTP), as previously described (12). For all subsequent experiments, the transepithelial electrical resistance and the equivalent current (Ieq) were recorded using a TECC-24 device as previously described (82). Briefly, to determine the baseline resistance, eight initial measurements were collected at ~2 min intervals. Changes in Ieq were determined after the addition of 6 μM benzamil (Benz), 10 μM FSK, 10 μM CFTR potentiator (Genistein (Gen) or VX-770 as indicated), and either 20 μM CFTRinh-172 or a mixture of 20 μM CFTRinh-172, 20 μM GlyH-101, and 20 μM Bumetanide (Inh mix) as indicated. TECC-24 assays were performed on days 21-32 after seeding.

### Statistical Analysis

All data were analyzed using ordinary linear models or linear mixed-effects models when multiple donors were studied with the donor as a random effect factor. The log transformation was performed before statistical testing when the variance differed greatly between groups as indicated in the figure legends. Analyses were performed with either the lm function in base R or the lmer function in the R packages *lme4* and *lmerTest*. Post hoc comparisons were performed using the general linear hypothesis test from the *multcomp* R package.

### Study Approval

Protocols for informed consent to obtain cells via nasal curettage and from explanted CF lungs were approved under the University of North Carolina’s Office of Human Research Ethics/Institutional Review Board studies 98-1015 and 03-1396, respectively.

## Author Contributions

REL, CAL, LH, and SHR conceived the project and designed the experiments with the assistance of JT and NC. The Bmi-1/hTERT lentiviral vector was prepared by JTM and LCM. Nasal curettage and review of clinical records were performed by MRK and AJK. Cell line generation and growth curve analyses were performed by REL, CAL, and LH. TMM, CAL, SCG, and REL performed whole-mount immunostaining and confocal imaging. Western blots and qRT-PCR was performed by LH. Electrophysiology experiments were performed by REL and SCG. REL, LH, JT, and NC assessed CC-90009’s effects on PTC readthrough. All of the statistical analysis was performed by HD. REL and CAL composed the manuscript and figures. All authors read, edited, and approved of the final version of the manuscript.

## Acknowledgements

The authors thank the cell donors; Dr. Snyder and the BioRepository & Precision Pathology Center Staff at Duke University; Drs. Blau, Mussaffi, and Prais at Schneider Children’s Hospital, Israel; and Dr. Kramer at Rabin Medical Center, Israel for facilitating the procurement of CF lungs. We thank Eric Roe for his help with reference formatting and the Marsico Lung Institute Tissue Procurement and Cell Culture Core for providing cells and media. NIH3T3J2 cells were kindly provided by Drs. Schlegel and Liu at Georgetown University. Novel CFTR modulators were provided by Dr. Lukacs at McGill University. Supported by Emily’s Entourage grant 17-5600; Cystic Fibrosis Foundation grants RANDEL17XX0, RANDEL20XX2, and BOUCHE19R0; and NIH grants 1T32GM133364, 1F31HL158197, 5KL2TR002490, 5R01HL071798, and 2P30DK065988.

## References

1. Cutting GR. Cystic fibrosis genetics: from molecular understanding to clinical application. Nat Rev Genet. 2015;16(1):45–56.

2. Ratjen F, et al. Cystic fibrosis. Nat Rev Dis Primers. 2015;1:15010.

3. Ramsey BW, et al. A CFTR potentiator in patients with cystic fibrosis and the G551D mutation. N Engl J Med. 2011;365(18):1663–1672.

4. Rowe SM, et al. Tezacaftor-ivacaftor in residual-function heterozygotes with cystic fibrosis. N Engl J Med. 2017;377(21):2024–2035.

5. Taylor-Cousar JL, et al. Tezacaftor-ivacaftor in patients with cystic fibrosis homozygous for Phe508del. N Engl J Med. 2017;377(21):2013–2023.

6. Wainwright CE, et al. Lumacaftor-ivacaftor in patients with cystic fibrosis homozygous for Phe508del CFTR. N Engl J Med. 2015;373(3):220–231.

7. Middleton PG, et al. Elexacaftor-tezacaftor-ivacaftor for cystic fibrosis with a single Phe508del allele. N Engl J Med. 2019;381(19):1809–1819.

8. Durmowicz AG, et al. The U.S. Food and Drug Administration’s experience with ivacaftor in cystic fibrosis. Establishing efficacy using in vitro data in lieu of a clinical trial. Ann Am Thorac Soc. 2018;15(1):1–2.

9. Van Goor F, et al. Effect of ivacaftor on CFTR forms with missense mutations associated with defects in protein processing or function. J Cyst Fibros. 2014;13(1):29–36.

10. Randell SH, et al. Primary epithelial cell models for cystic fibrosis research. Methods Mol Biol. 2011;742:285–310.

11. Van Goor F, et al. Rescue of DeltaF508-CFTR trafficking and gating in human cystic fibrosis airway primary cultures by small molecules. Am J Physiol Lung Cell Mol Physiol. 2006;290(6):L1117–1130.

12. Gentzsch M, et al. Pharmacological rescue of conditionally reprogrammed cystic fibrosis bronchial epithelial cells. Am J Respir Cell Mol Biol. 2017;56(5):568–574.

13. Brewington JJ, et al. Brushed nasal epithelial cells are a surrogate for bronchial epithelial CFTR studies. JCI Insight. 2018;3(13).

14. Mosler K, et al. Feasibility of nasal epithelial brushing for the study of airway epithelial functions in CF infants. J Cyst Fibros. 2008;7(1):44–53.

15. Fulcher ML, et al. Novel human bronchial epithelial cell lines for cystic fibrosis research. Am J Physiol Lung Cell Mol Physiol. 2009;296(1):L82–91.

16. Munye MM, et al. BMI-1 extends proliferative potential of human bronchial epithelial cells while retaining their mucociliary differentiation capacity. Am J Physiol Lung Cell Mol Physiol. 2017;312(2):L258–l267.

17. Yip YL, et al. Efficient immortalization of primary nasopharyngeal epithelial cells for EBV infection study. PLoS One. 2013;8(10):e78395.

18. Baradaran-Heravi A, et al. Effect of small molecule eRF3 degraders on premature termination codon readthrough. Nucleic Acids Res. 2021;49(7):3692–3708.

19. Awatade NT, et al. Human primary epithelial cell models: promising tools in the era of cystic fibrosis personalized medicine. Front Pharmacol. 2018;9:1429.

20. Fulcher ML, Randell SH. Human nasal and tracheo-bronchial respiratory epithelial cell culture. Methods Mol Biol. 2013;945:109–121.

21. Reynolds SD, et al. Airway progenitor clone formation is enhanced by Y-27632-dependent changes in the transcriptome. Am J Respir Cell Mol Biol. 2016;55(3):323–336.

22. Wolf S, et al. Conditional reprogramming of pediatric airway epithelial cells: a new human model to investigate early-life respiratory disorders. Pediatr Allergy Immunol. 2017;28(8):810–817.

23. Kmit A, et al. Extent of rescue of F508del-CFTR function by VX-809 and VX-770 in human nasal epithelial cells correlates with SNP rs7512462 in SLC26A9 gene in F508del/F508del Cystic Fibrosis patients. Biochim Biophys Acta Mol Basis Dis. 2019;1865(6): 1323–1331.

24. Kreft ME, et al. Different culture conditions affect drug transporter gene expression, ultrastructure, and permeability of primary human nasal epithelial cells. Pharm Res. 2020;37(9):170.

25. Shao D, et al. Optimization of human nasal epithelium primary culture conditions for optimal proton oligopeptide and organic cation transporters expression in vitro. Int J Pharm. 2013;441(1-2):334–342.

26. Fulcher ML, et al. Well-differentiated human airway epithelial cell cultures. Methods Mol Med. 2005;107:183–206.

27. Neuberger T, et al. Use of primary cultures of human bronchial epithelial cells isolated from cystic fibrosis patients for the pre-clinical testing of CFTR modulators. Methods Mol Biol. 2011;741:39–54.

28. Knowles MR, et al. Activation by extracellular nucleotides of chloride secretion in the airway epithelia of patients with cystic fibrosis. N Engl J Med. 1991;325(8):533–538.

29. Veit G, et al. Structure-guided combination therapy to potently improve the function of mutant CFTRs. Nat Med. 2018;24(11):1732–1742.

30. Keating D, et al. VX-445-tezacaftor-ivacaftor in patients with cystic fibrosis and one or two Phe508del alleles. N Engl J Med. 2018;379(17):1612–1620.

31. Valley HC, et al. Isogenic cell models of cystic fibrosis-causing variants in natively expressing pulmonary epithelial cells. J Cyst Fibros. 2019;18(4):476–483.

32. Mutyam V, et al. Therapeutic benefit observed with the CFTR potentiator, ivacaftor, in a CF patient homozygous for the W1282X CFTR nonsense mutation. J Cyst Fibros. 2017;16(1):24–29.

33. Shalev M, Baasov T. When proteins start to make sense: fine-tuning aminoglycosides for PTC suppression therapy. Medchemcomm. 2014;5(8):1092–1105.

34. Konstan MW, et al. Efficacy and safety of ataluren in patients with nonsense-mutation cystic fibrosis not receiving chronic inhaled aminoglycosides: The international, randomized, double-blind, placebo-controlled Ataluren Confirmatory Trial in Cystic Fibrosis (ACT CF). J Cyst Fibros. 2020;19(4):595–601.

35. Kerem E, et al. Ataluren for the treatment of nonsense-mutation cystic fibrosis: a randomised, double-blind, placebo-controlled phase 3 trial. Lancet Respir Med. 2014;2(7):539–547.

36. Linde L, et al. Nonsense-mediated mRNA decay affects nonsense transcript levels and governs response of cystic fibrosis patients to gentamicin. J Clin Invest. 2007;117(3):683–692.

37. Aksit MA, et al. Decreased mRNA and protein stability of W1282X limits response to modulator therapy. J Cyst Fibros. 2019;18(5):606–613.

38. Laselva O, et al. Functional rescue of c.3846G>A (W1282X) in patient-derived nasal cultures achieved by inhibition of nonsense mediated decay and protein modulators with complementary mechanisms of action. J Cyst Fibros. 2020;19(5):717–727.

39. Haggie PM, et al. Correctors and potentiators rescue function of the truncated W1282X-cystic fibrosis transmembrane regulator (CFTR) translation product. J Biol Chem. 2017;292(3):771–785.

40. Wang W, et al. Robust stimulation of W1282X-CFTR channel activity by a combination of allosteric modulators. PLoS One. 2016;11(3):e0152232.

41. Canessa CM, et al. Amiloride-sensitive epithelial Na+ channel is made of three homologous subunits. Nature. 1994;367(6462):463–467.

42. Kimura T, et al. Deletion of the ubiquitin ligase Nedd4L in lung epithelia causes cystic fibrosis-like disease. Proc Natl Acad Sci U S A. 2011;108(8):3216–3221.

43. Surka C, et al. CC-90009, a novel cereblon E3 ligase modulator, targets acute myeloid leukemia blasts and leukemia stem cells. Blood. 2021;137(5):661–677.

44. Shei RJ, et al. The epithelial sodium channel (ENaC) as a therapeutic target for cystic fibrosis. Curr Opin Pharmacol. 2018;43:152–165.

45. Van Goor F, et al. Rescue of CF airway epithelial cell function in vitro by a CFTR potentiator, VX-770. Proc Natl Acad Sci U S A. 2009;106(44):18825–18830.

46. Van Goor F, et al. Correction of the F508del-CFTR protein processing defect in vitro by the investigational drug VX-809. Proc Natl Acad Sci U S A. 2011;108(46):18843–18848.

47. De Boeck K, et al. Efficacy and safety of ivacaftor in patients with cystic fibrosis and a non-G551D gating mutation. J Cyst Fibros. 2014;13(6):674–680.

48. Donaldson SH, et al. Tezacaftor/ivacaftor in subjects with cystic fibrosis and F508del/F508del-CFTR or F508del/G551D-CFTR. Am J Respir Crit Care Med. 2018;197(2):214–224.

49. Suprynowicz FA, et al. Conditionally reprogrammed cells represent a stem-like state of adult epithelial cells. Proc Natl Acad Sci U S A. 2012;109(49):20035–20040.

50. Wilschanski M, et al. A pilot study of the effect of gentamicin on nasal potential difference measurements in cystic fibrosis patients carrying stop mutations. Am J Respir Crit Care Med. 2000;161(3 Pt 1):860–865.

51. Wilschanski M, et al. Gentamicin-induced correction of CFTR function in patients with cystic fibrosis and CFTR stop mutations. N Engl J Med. 2003;349(15):1433–1441.

52. Dabrowski M, et al. Advances in therapeutic use of a drug-stimulated translational readthrough of premature termination codons. Mol Med. 2018;24(1):25.

53. Nagel-Wolfrum K, et al. Targeting nonsense mutations in diseases with translational read-through-inducing drugs (TRIDs). BioDrugs. 2016;30(2):49–74.

54. Keeling KM, et al. Therapeutics based on stop codon readthrough. Annu Rev Genomics Hum Genet. 2014;15:371–394.

55. Xie J, et al. New developments in aminoglycoside therapy and ototoxicity. Hear Res. 2011;281(1-2):28–37.

56. Zhouravleva G, et al. Termination of translation in eukaryotes is governed by two interacting polypeptide chain release factors, eRF1 and eRF3. EMBO J. 1995;14(16):4065–4072.

57. Frolova L, et al. Eukaryotic polypeptide chain release factor eRF3 is an eRF1- and ribosome-dependent guanosine triphosphatase. RNA. 1996;2(4):334–341.

58. Carnes J, et al. Stop codon suppression via inhibition of eRF1 expression. RNA. 2003;9(6):648–653.

59. Chauvin C, et al. Involvement of human release factors eRF3a and eRF3b in translation termination and regulation of the termination complex formation. Mol Cell Biol. 2005;25(14):5801–5811.

60. Huang L, et al. Targeting translation termination machinery with antisense oligonucleotides for diseases caused by nonsense mutations. Nucleic Acid Ther. 2019;29(4):175–186.

61. Sharma J, et al. A small molecule that induces translational readthrough of CFTR nonsense mutations by eRF1 depletion. Nat Commun. 2021;12(1):4358.

62. Matyskiela ME, et al. A novel cereblon modulator recruits GSPT1 to the CRL4(CRBN) ubiquitin ligase. Nature. 2016;535(7611):252–257.

63. Lu G, et al. Elucidating the mechanism of action of CC-90009, a novel cereblon E3 ligase modulator, in AML via genome-wide CRISPR screen. Blood. 2019;134(Supplement_1):405–405.

64. Lopez-Girona A, et al. CC-90009, a novel cereblon E3 ligase modulator, targets GSPT1 for degradation to induce potent tumoricidal activity against acute myeloid leukemia (AML). Blood. 2019;134:2703.

65. Uy GL, et al. Clinical activity of CC-90009, a cereblon E3 ligase modulator and first-in-class GSPT1 degrader, as a single agent in patients with relapsed or refractory acute myeloid leukemia (R/R AML): First results from a phase I dose-finding study. Blood. 2019;134(Supplement_1):232–232.

66. Fan J, et al. Pharmacodynamic responses to CC-90009, a novel cereblon E3 ligase modulator, in a phase I dose-escalation study in relapsed or refractory acute myeloid leukemia (R/R AML). Blood. 2019;134(Supplement_1):2547–2547.

67. Jin L, et al. A novel cereblon E3 ligase modulator eradicates acute myeloid leukemia stem cells through degradation of translation termination factor GSPT1. Blood. 2019;134:3940.

68. Lee RE, et al. Assessing human airway epithelial progenitor cells for cystic fibrosis cell therapy. Am J Respir Cell Mol Biol. 2020;63(3):374–385.

69. Bidou L, et al. Sense from nonsense: therapies for premature stop codon diseases. Trends Mol Med. 2012;18(11):679–688.

70. Keeling KM, et al. Suppression of premature termination codons as a therapeutic approach. Crit Rev Biochem Mol Biol. 2012;47(5):444–463.

71. Aliouat A, et al. Divergent effects of translation termination factor eRF3A and nonsense-mediated mRNA decay factor UPF1 on the expression of uORF carrying mRNAs and ribosome protein genes. RNA Biol. 2020;17(2):227–239.

72. Pratt JH. Central role for ENaC in development of hypertension. J Am Soc Nephrol. 2005;16(11):3154–3159.

73. Song J, et al. Regulation of blood pressure, the epithelial sodium channel (ENaC), and other key renal sodium transporters by chronic insulin infusion in rats. Am J Physiol Renal Physiol. 2006;290(5):F1055–1064.

74. Abriel H, et al. Defective regulation of the epithelial Na+ channel by Nedd4 in Liddle’s syndrome. J Clin Invest. 1999;103(5):667–673.

75. Shimkets RA, et al. Liddle’s syndrome: heritable human hypertension caused by mutations in the beta subunit of the epithelial sodium channel. Cell. 1994;79(3):407–414.

76. Chang SS, et al. Mutations in subunits of the epithelial sodium channel cause salt wasting with hyperkalaemic acidosis, pseudohypoaldosteronism type 1. Nat Genet. 1996;12(3):248–253.

77. Welch EM, et al. PTC124 targets genetic disorders caused by nonsense mutations. Nature. 2007;447(7140):87–91.

78. Liu X, et al. Conditional reprogramming and long-term expansion of normal and tumor cells from human biospecimens. Nat Protoc. 2017;12(2):439–451.

79. Fu B, et al. Keratinocyte growth conditions modulate telomerase expression, senescence, and immortalization by human papillomavirus type 16 E6 and E7 oncogenes. Cancer Res. 2003;63(22):7815–7824.

80. Liu X, et al. HPV E7 contributes to the telomerase activity of immortalized and tumorigenic cells and augments E6-induced hTERT promoter function. Virology. 2008;375(2):611–623.

81. Ghosh A, et al. Chronic e-cigarette exposure alters the human bronchial epithelial proteome. Am J Respir Crit Care Med. 2018;198(1):67–76.

82. Liang F, et al. High-throughput screening for readthrough modulators of CFTR PTC mutations. SLAS Technol. 2017;22(3):315–324.

