## Supplemental Figure 1 for "Robust W1282X-CFTR rescue by a small molecule GSPT1 degrader"

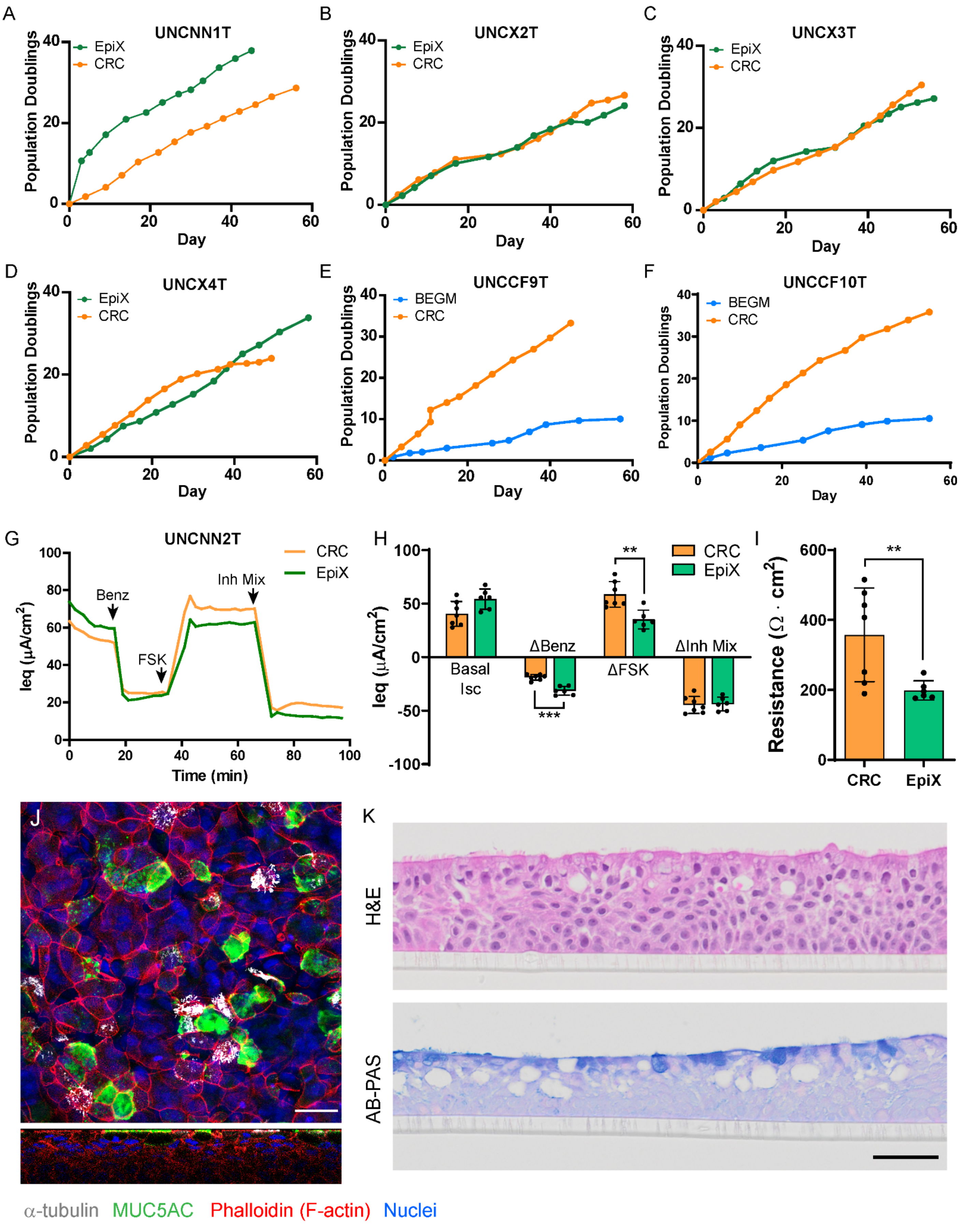
A
UNCNN1T
B
UNCX2T
C
UNCX3T
D
UNCX4T
E
UNCCF9T
F
UNCCF10T
EpiX
CRC
BEGM
Population Doublings
Day
G
UNCNN2T
Benz
FSK
Inh Mix
Ieq (μA/cm²)
Time (min)
H
Basal Isc
ΔBenz
ΔFSK
ΔInh Mix
I
Resistance (Ω · cm²)
J
K
H&E
AB-PAS
α-tubulin MUC5AC Phalloidin (F-actin) Nuclei

*Supplemental Figure 1. CRC and EpiX medium enable robust nasal cell expansion.* A-D) Bmi-1/hTERT nasal cell line population doublings over time in CRC and EpiX media. A) UNCNN1T; B) UNCX2T; C) UNCX3T; D) UNCX4T. E-F) Bmi-1/hTERT bronchial cell line population doublings over time in CRC and bronchial epithelial growth media (BEGM). E) UNCCF9T; F) UNCCF10T. G) Representative TECC-24 tracing of UNCNN2T expanded in CRC or EpiX media and differentiated in Pneumacult ALI media. H) Basal Ieq and ΔIeq in response to Benzamil, FSK, and Inh Mix in UNCNN1T and UNCNN2T expanded in CRC or EpiX media. N=2-4 for each donor. Data were analyzed using a linear mixed-effects model with the donor as a random effect factor. I) Baseline resistance values. N=2-4 for each donor. Data were analyzed using a linear mixed-effects model with the donor as a random effect factor. J) Whole-mount immunostaining of UNCNN2T P6 expanded in EpiX. α-tubulin (white), MUC5AC (green), Phalloidin (F-actin, red), and Hoechst (Nuclei, blue). Scale bar = 25 μm. K) H&E and AB-PAS staining of UNCNN2T P6 expanded in EpiX. Scale bar = 50 μm. All data presented as mean ± SD. ** = $p<0.01$; *** = $p<0.001$.

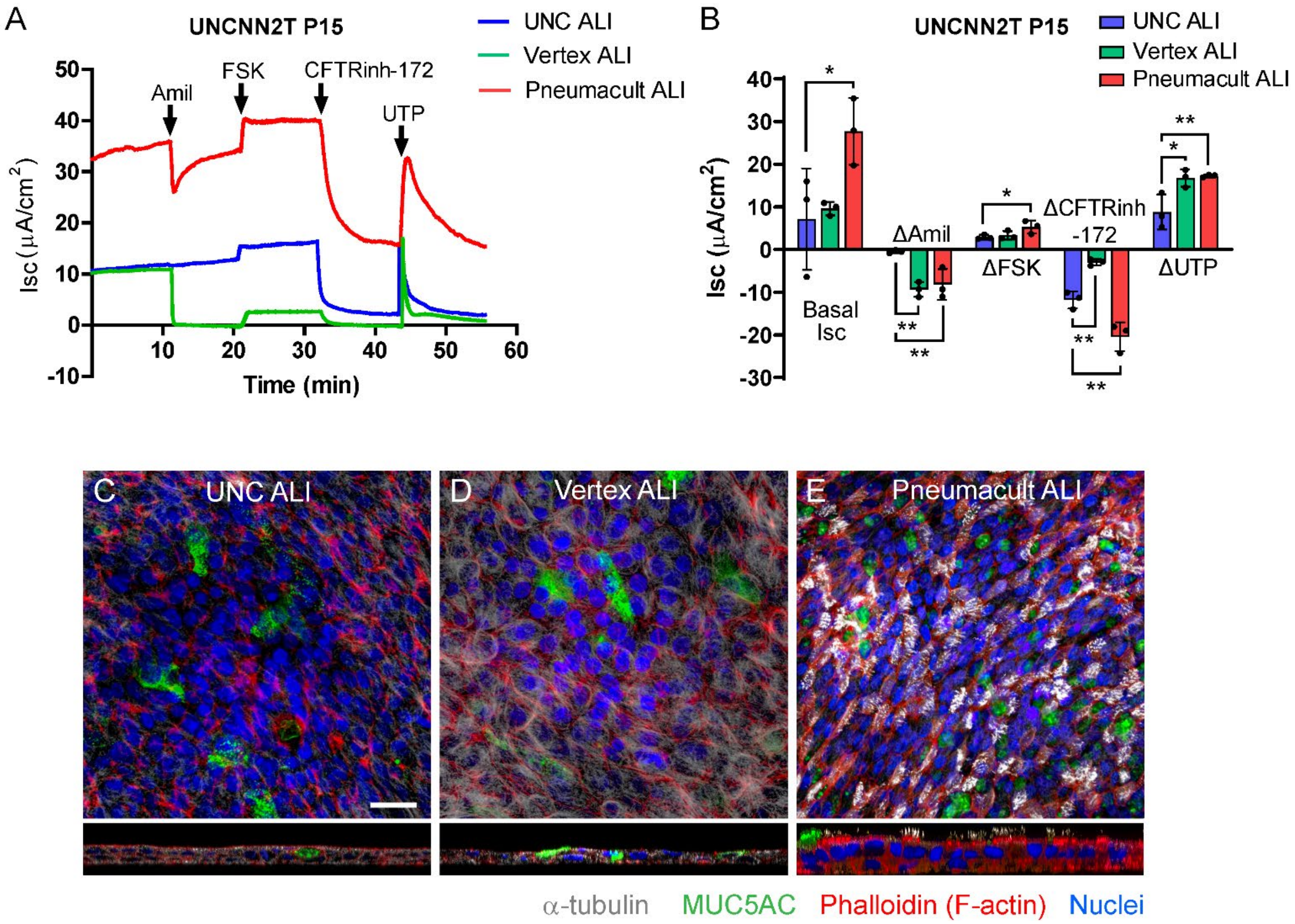


*Supplemental Figure 2. Pneumacult ALI promotes healthy differentiation of Bmi-1/hTERT nasal cell lines.* A-B) Ussing chamber measurements of UNCNN2T P15 cells differentiated with UNC ALI, Vertex ALI, or Pneumacult ALI media. A) Representative Ussing tracing. B) Change in short-circuit current (ΔIsc) in response to Amiloride (Amil), FSK, CFTRinh-172, and Uridine-5'-triphosphate (UTP). N=3. Data were analyzed using an ordinary linear model. C-E) Whole-mount immunostaining of UNCNN2T P15 cells differentiated in UNC ALI (C), Vertex ALI (D), or Pneumacult ALI (E). α-tubulin (white), MUC5AC (green), Phalloidin (F-actin, red), and Hoechst (Nuclei, blue). Scale bar = 25 μm. All data presented as mean ± SD. * = $p<0.05$; ** = $p<0.01$.

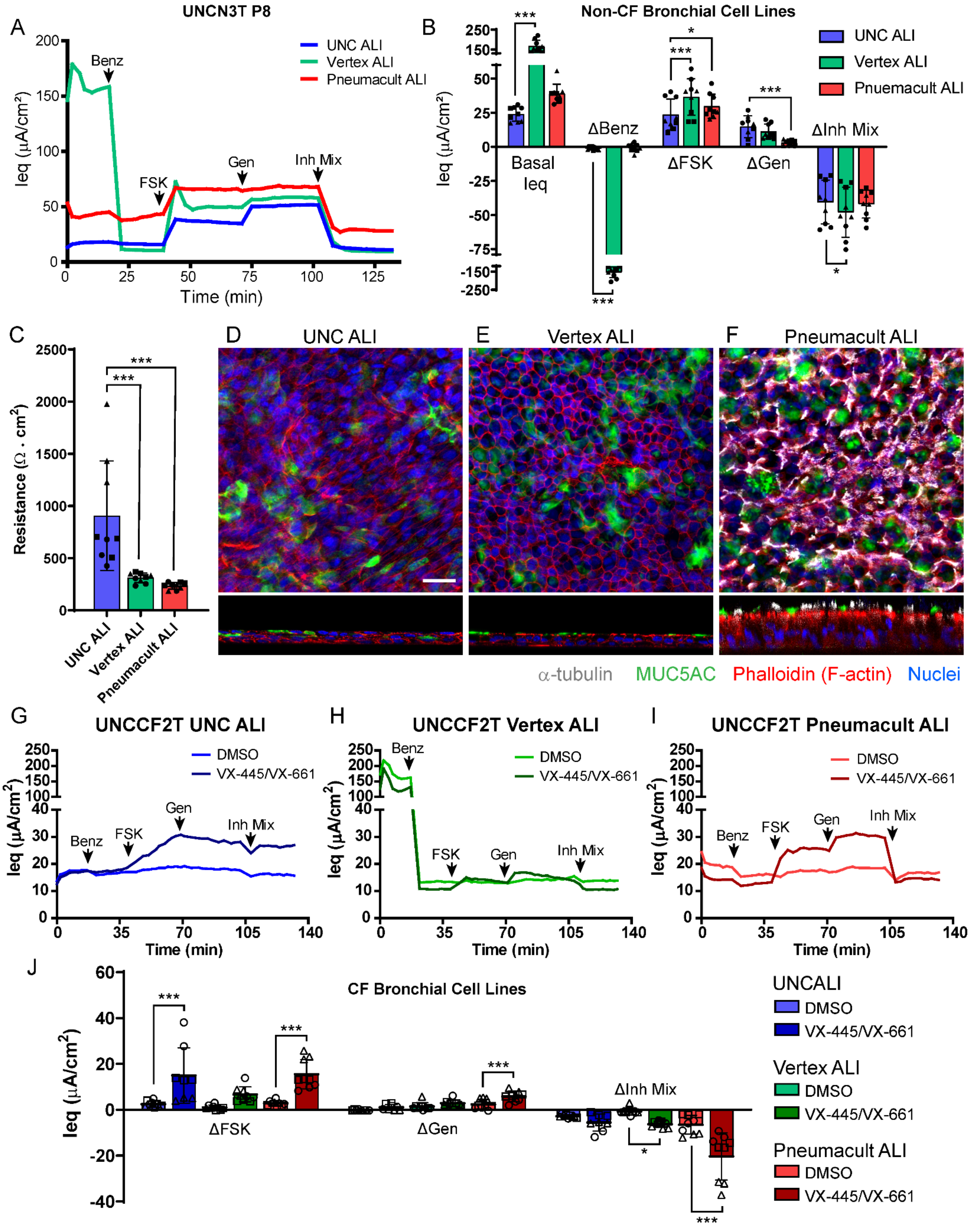

A
UNCN3T P8
Benz
FSK
Gen
Inh Mix
UNC ALI
Vertex ALI
Pneumacult ALI
Ieq (μA/cm²)
Time (min)
B
Non-CF Bronchial Cell Lines
UNC ALI
Vertex ALI
Pnuemacult ALI
Basal Ieq
ΔBenz
ΔFSK
ΔGen
ΔInh Mix
C
Resistance (Ω · cm²)
UNC ALI
Vertex ALI
Pneumacult ALI
D
UNC ALI
E
Vertex ALI
F
Pneumacult ALI
α-tubulin
MUC5AC
Phalloidin (F-actin)
Nuclei
G
UNCCF2T UNC ALI
H
UNCCF2T Vertex ALI
I
UNCCF2T Pneumacult ALI
DMSO
VX-445/VX-661
J
CF Bronchial Cell Lines
ΔFSK
ΔGen
ΔInh Mix
UNCALI
Vertex ALI
Pneumacult ALI

*Supplemental Figure 3. Pneumacult ALI promotes differentiation of CF and non-CF Bmi-1/hTERT bronchial cell lines.* A-C) TECC-24 measurements of three non-CF Bmi-1/hTERT bronchial cell lines previously published by our lab, UNCN1T P8, UNCN2T P9, and UNCN3T P8, differentiated with UNC ALI, Vertex ALI, or Pneumacult ALI media. A) Representative tracing of UNCN3T P8 grown in the three differentiation conditions. B) Basal Ieq and ΔIeq in response to Benz, FSK, Gen, and Inh Mix. Data were analyzed using a linear mixed-effects model with the donor as a random effect factor. C) Baseline resistance values. Log transformed data were analyzed using a linear mixed-effects model with the donor as a random effect factor. B-C) UNCN1T displayed as circles, UNCN2T displayed as squares, and UNCN3T displayed as triangles. N=3 for each cell line. D-F) Whole-mount immunostaining of UNCN3T P8 differentiated with UNC ALI (D), Vertex ALI (E), or Pneumacult ALI (F). α-tubulin (white), MUC5AC (green), Phalloidin (F-actin, red), and Hoechst (Nuclei, blue). Scale bar = 25 µm. G-J) TECC-24 measurements from three CF bronchial cell lines generated from F508del/F508del donors, UNCCF1T P8, UNCCF2T P7, and UNCCF3T P8, pretreated with VX-445/VX-661 (both at 5 µM) or DMSO for 48 hours. G-I) Representative tracings of UNCCF2T grown in UNC ALI (G), Vertex ALI (H), or Pneumacult ALI (I). J) ΔIeq in response to FSK, Gen, and Inh Mix. UNCCF1T displayed as circles, UNCCF2T displayed as squares, and UNCCF3T displayed as triangles. N=2-3 for each cell line. Data were analyzed using a linear mixed-effects model with the donor as a random effect factor. All data presented as mean ± SD. * = $p<0.05$; *** = $p<0.001$.

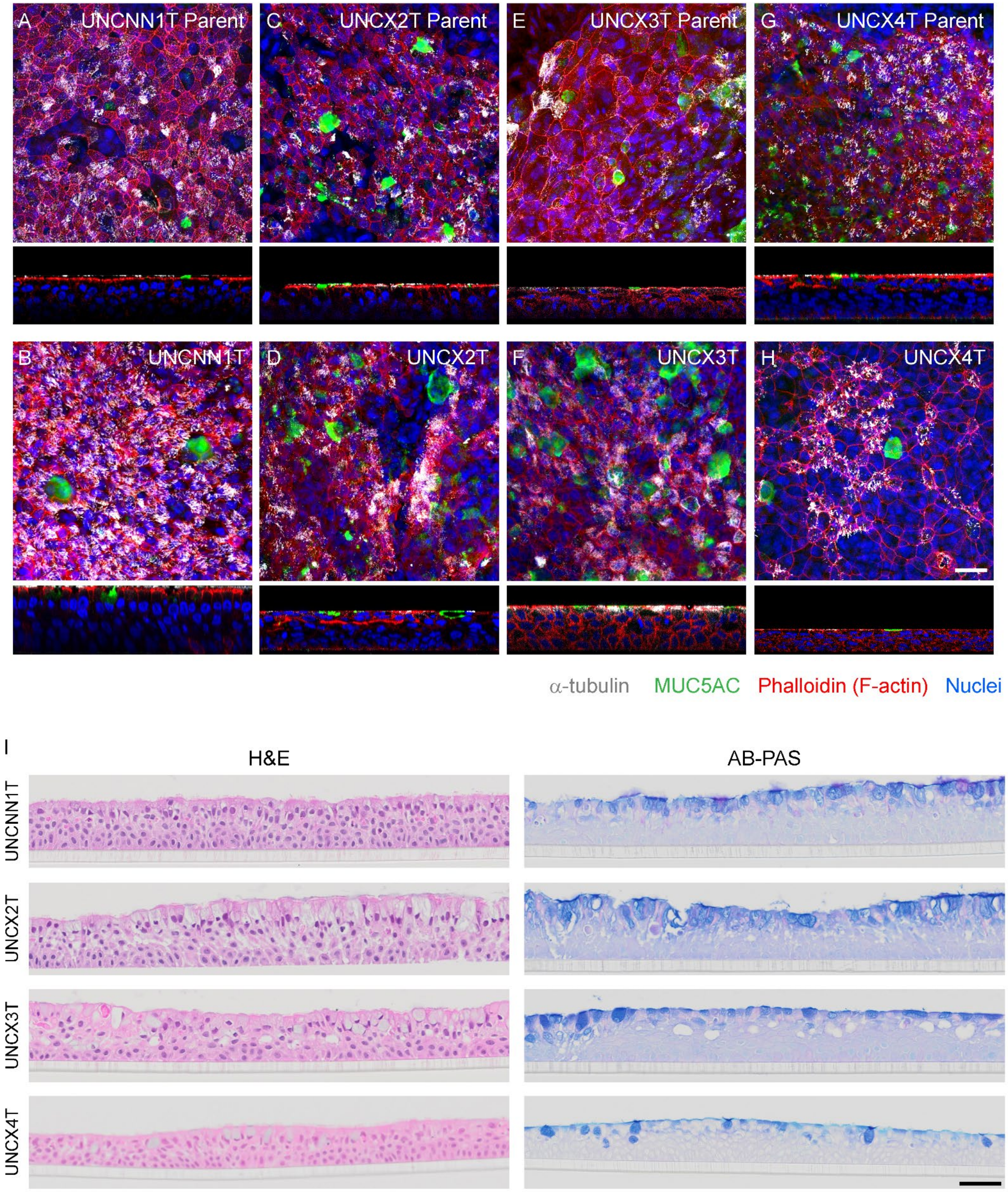


*Supplemental Figure 4. Morphology of nasal cell lines.* A-H) Whole-mount immunostaining of P2 parent cells and P6 cell line for UNCNN1T (A-B), UNCX2T (C-D), UNCX3T (E-F), and UNCX4T (G-H). α-tubulin (white), MUC5AC (green), Phalloidin (F-actin, red), and Hoechst (Nuclei, blue). Scale bar = 25 μm. I) H&E and AB-PAS staining of UNCNN1T, UNCX2T, UNCX3T, and UNCX4T at P5-7. Scale bar = 50 μm.

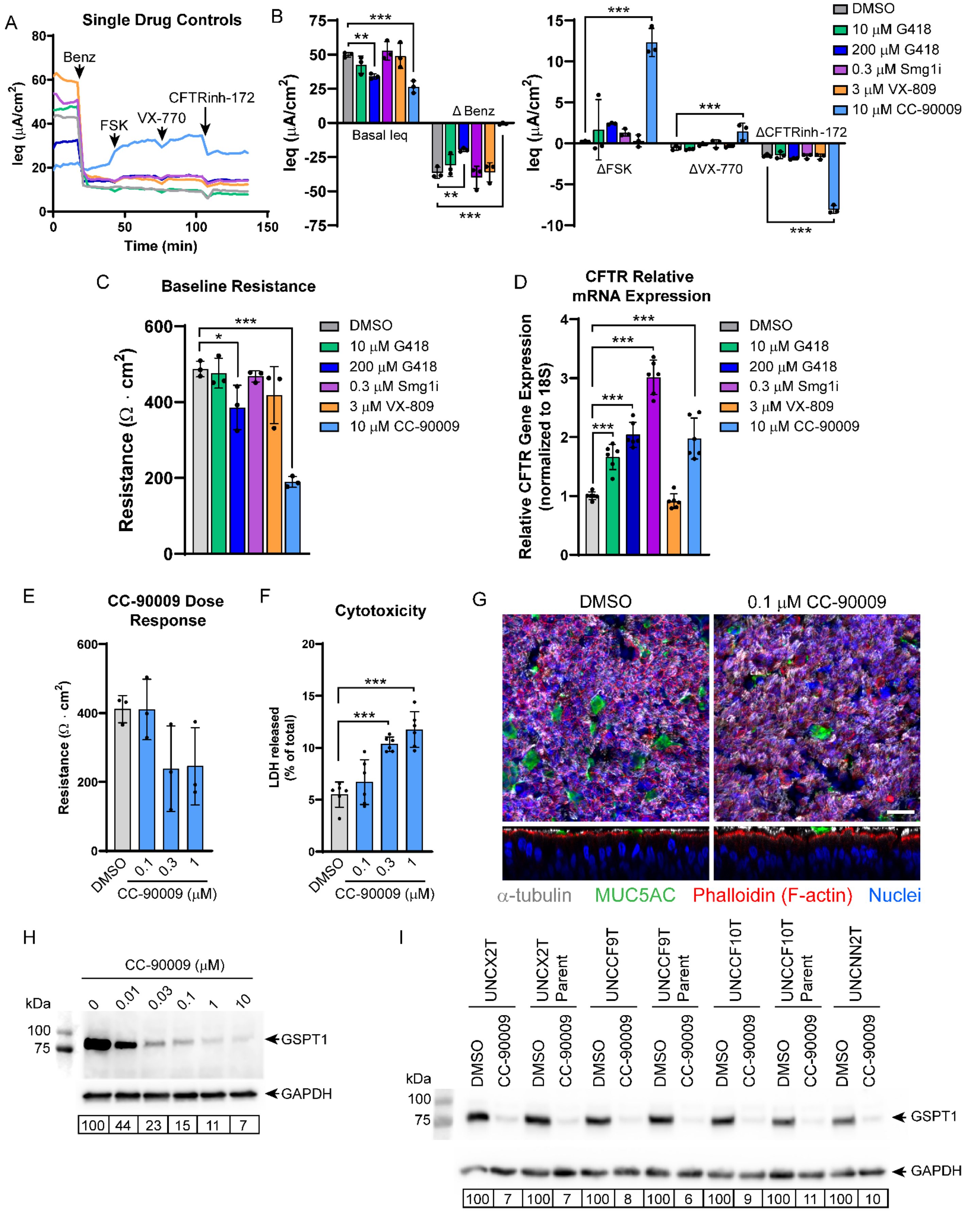
A
Single Drug Controls
Benz
FSK
VX-770
CFTRinh-172
Ieq (μA/cm²)
Time (min)
B
Basal Ieq
Δ Benz
ΔFSK
ΔVX-770
ΔCFTRinh-172
Ieq (μA/cm²)
DMSO
10 μM G418
200 μM G418
0.3 μM Smg1i
3 μM VX-809
10 μM CC-90009
C
Baseline Resistance
Resistance (Ω · cm²)
D
CFTR Relative mRNA Expression
Relative CFTR Gene Expression (normalized to 18S)
E
CC-90009 Dose Response
Resistance (Ω · cm²)
DMSO
CC-90009 (μM)
F
Cytotoxicity
LDH released (% of total)
G
DMSO
0.1 μM CC-90009
α-tubulin
MUC5AC
Phalloidin (F-actin)
Nuclei
H
CC-90009 (μM)
kDa
GSPT1
GAPDH
100 44 23 15 11 7
I
UNCX2T
UNCX2T Parent
UNCCF9T
UNCCF9T Parent
UNCCF10T
UNCCF10T Parent
UNCNN2T
DMSO
CC-90009
GSPT1
GAPDH
100 7 100 7 100 8 100 6 100 9 100 11 100 10

*Supplemental Figure 5. CC-90009 rescues W1282X CFTR without aminoglycosides or NMD inhibition.* A-C) TECC-24 measurements of the UNCX2T cell line pretreated with DMSO, 10 μM or 200 μM G418, 0.3 μM SMG1i, 3 μM VX-809, or 10 μM CC-90009 for 48 hours. A) Representative TECC-24 tracing. B) Basal Ieq and ΔIeq in response to Benz (left). ΔIeq in response to FSK, VX-770, and CFTRinh-172 (right). N=3. C) Baseline resistance values. N=3. D) Relative CFTR mRNA expression by qRT-PCR in UNCX2T pretreated with DMSO, 10 μM or 200 μM G418, 0.3 μM SMG1i, 3 μM VX-809, or 10 μM CC-90009 for 48 hours. N=6. E-F) Dose response of CC-90009 in UNCX2T parent cells (24-hour treatment). E) Transepithelial resistance. N=3. F) Cytotoxicity measured by LDH release. N=3. G) Whole-mount immunostaining of UNCCF10T cells pretreated with DMSO or 0.1 μM CC-90009 for 24 hours. α-tubulin (white), MUC5AC (green), Phalloidin (F-actin, red), and Hoechst (Nuclei, blue). Scale bar = 25 μm. H) Western blot for GSPT1 in UNCCF9T parent cells pretreated with escalating doses of CC-90009 for 24 hours. Protein expression normalized to GAPDH and relative to the DMSO control quantified below. I) Western blot for GSPT1 in all W1282X cell lines and parent cells and the UNCNN2T cell line pretreated with DMSO or 0.1 μM CC-90009 for 24 hours. Protein expression normalized to GAPDH and relative to the paired DMSO control quantified below. All data were analyzed using ordinary linear models and presented as mean ± SD. * = $p<0.05$; ** = $p<0.01$; *** = $p<0.001$.

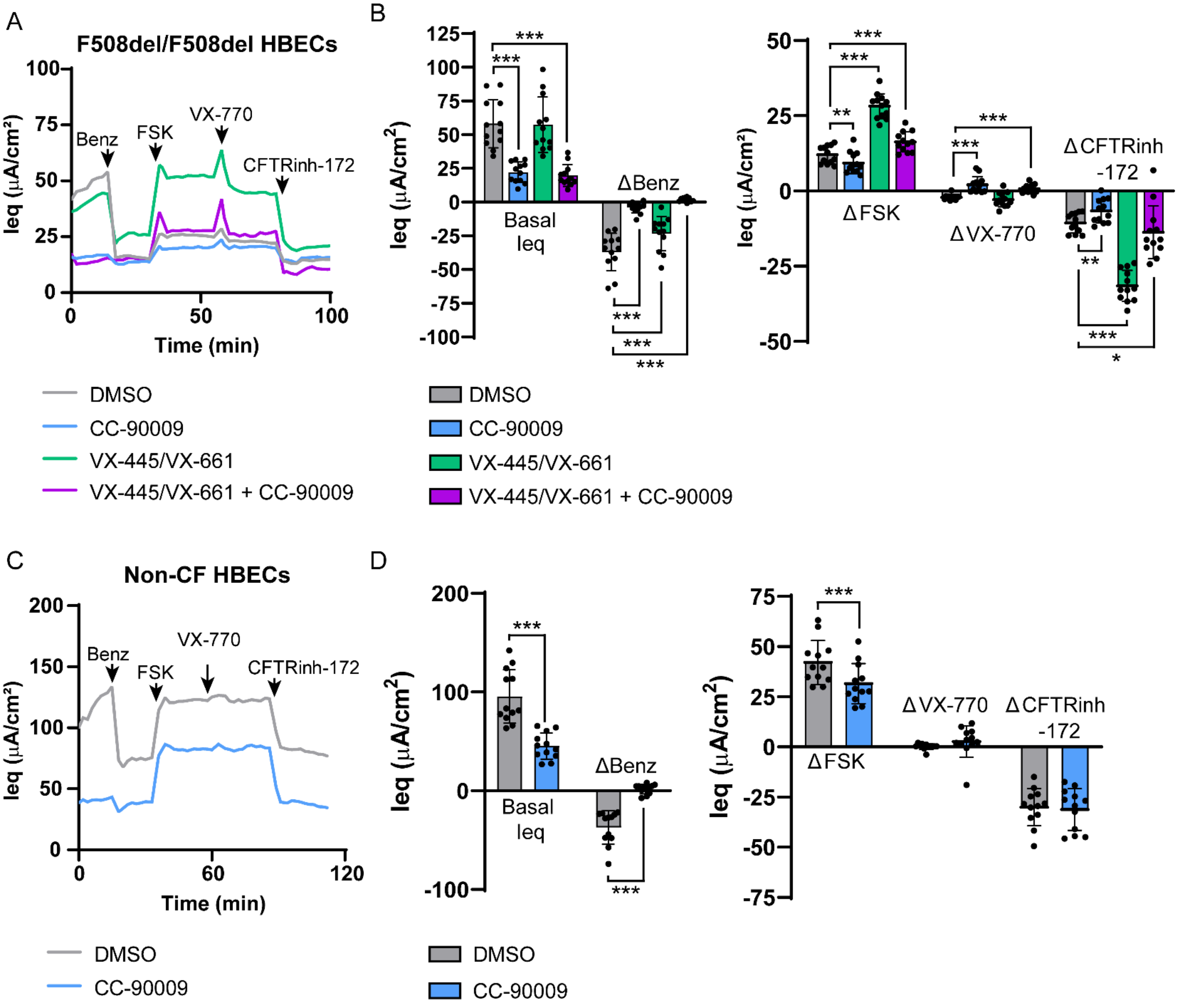


*Supplemental Figure 6. CC-90009 does not improve CFTR function in F508del homozygous or non-CF primary bronchial cells.* A-B) TECC-24 measurements of F508del/F508del HBECs pretreated with DMSO, 0.1 μM CC-90009, VX-445/VX-661 (both at 5 μM), or VX-445/VX-661(both at 5 μM) + 0.1 μM CC-90009 for 48 hours. A) Representative TECC-24 tracing. B) Basal Ieq and ΔIeq in response to Benz (left). ΔIeq in response to FSK, VX-770, and CFTRinh-172 (right). Biological N=4; 3 replicates per donor. C-D) TECC-24 measurements of non-CF primary HBECs pretreated with DMSO or 0.1 μM CC-90009 for 24 hours. C) Representative TECC-24 tracing. D) Basal Ieq and ΔIeq in response to Benz (left). ΔIeq in response to FSK, VX-770, and CFTRinh-172 (right). Biological N = 4; 3 replicates per donor. All data were analyzed using linear mixed-effects models with the donor as a random effect factor. Data represented as mean ± SD. * = $p<0.05$; ** = $p<0.01$; *** = $p<0.001$.
